# Dual inhibition of ATR and DNA-PKcs radiosensitizes ATM-mutant prostate cancer

**DOI:** 10.1101/2024.07.10.602941

**Authors:** Mia Hofstad, Andrea Woods, Karla Parra, Zoi E. Sychev, Alice Mazzagatti, Lan Yu, Collin Gilbreath, Peter Ly, Justin M. Drake, Ralf Kittler

## Abstract

In advanced castration resistant prostate cancer (CRPC), mutations in the DNA damage response (DDR) gene *ataxia telangiectasia mutated* (*ATM*) are common. While poly(ADP-ribose) polymerase inhibitors are approved in this context, their clinical efficacy remains limited. Thus, there is a compelling need to identify alternative therapeutic avenues for *ATM* mutant prostate cancer patients. Here, we generated matched ATM-proficient and ATM-deficient CRPC lines to elucidate the impact of ATM loss on DDR in response to DNA damage via irradiation. Through unbiased phosphoproteomic screening, we unveiled that ATM-deficient CRPC lines maintain dependence on downstream ATM targets through activation of ATR and DNA-PKcs kinases. Dual inhibition of ATR and DNA-PKcs effectively inhibited downstream γH2AX foci formation in response to irradiation and radiosensitized ATM-deficient lines to a greater extent than either ATM-proficient controls or single drug treatment. Further, dual inhibition abrogated residual downstream ATM pathway signaling and impaired replication fork dynamics. To circumvent potential toxicity, we leveraged the RUVBL1/2 ATPase inhibitor Compound B, which leads to the degradation of both ATR and DNA-PKcs kinases. Compound B effectively radiosensitized ATM-deficient CRPC *in vitro* and *in vivo*, and impacted replication fork dynamics. Overall, dual targeting of both ATR and DNA-PKcs is necessary to block DDR in ATM-deficient CRPC, and Compound B could be utilized as a novel therapy in combination with irradiation in these patients.

**Graphical Abstract:** 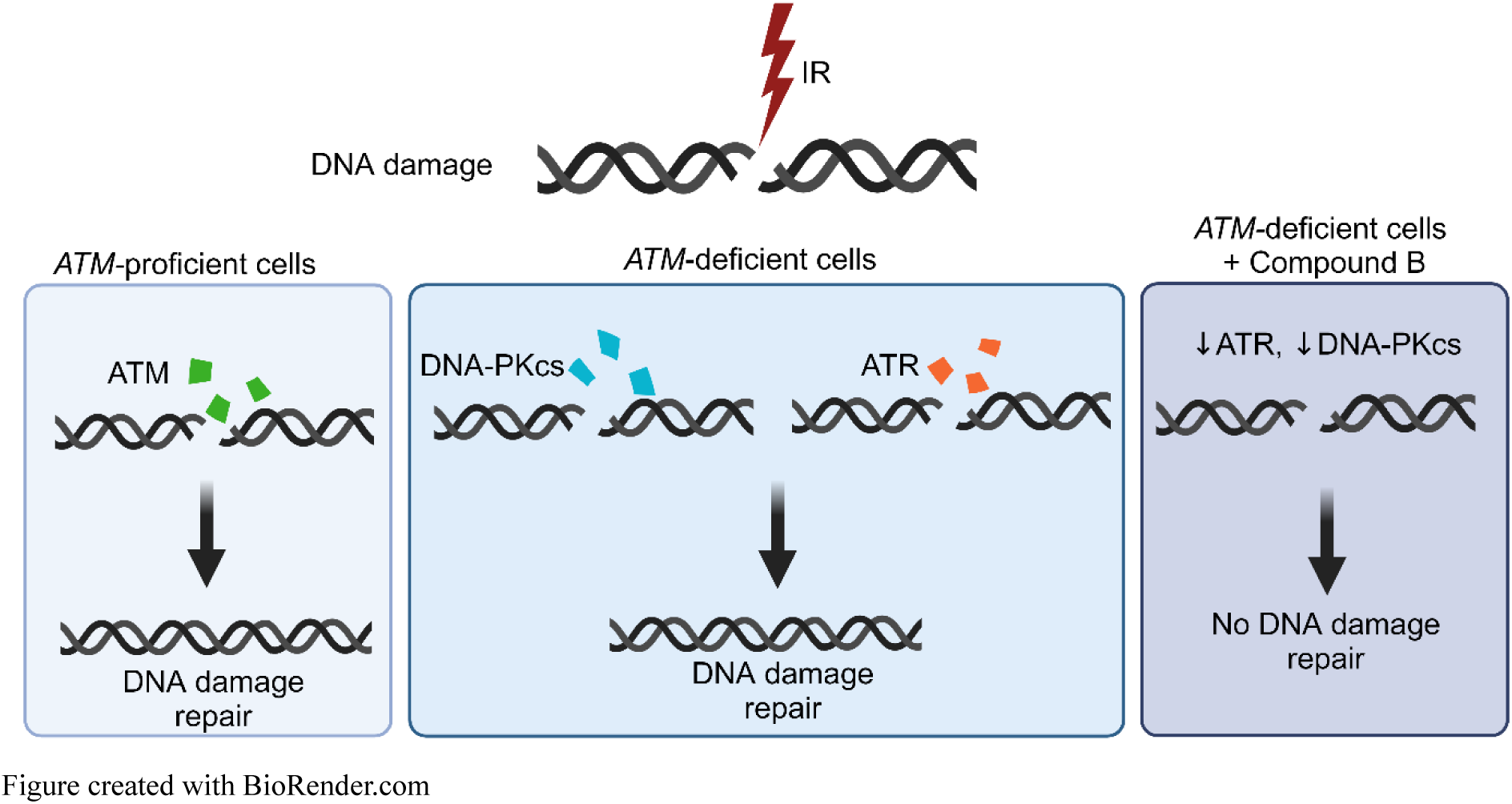

## Introduction

Prostate cancer is a leading cause of morbidity and mortality for men, accounting for ∼288,000 new cases and 11% of estimated cancer-related deaths in the United States in 2023 [1]. The vast majority of newly diagnosed cases are localized and are successfully managed with either active surveillance, radical prostatectomy, and/or targeted radiation [2]. However, a subset of patients initially present with metastases or progress on local therapy, requiring androgen deprivation therapy (ADT). While patients initially demonstrate a robust response to ADT, many ultimately progress to castration-resistant prostate cancer (CRPC) [3]. Much progress has been made in targeting the continued androgen dependency of CRPCs with the advent of the second-generation anti-androgens such as abiraterone acetate and enzalutamide [4–6]. However, patients commonly develop resistance to these second-generation anti-androgens, underscoring the need for alternative targeted modalities [7, 8].

Germline and somatic loss of function mutations in DNA damage response (DDR) genes, essential for homologous recombination (HR), have recently emerged as oncogenotypes with synthetic lethality to poly-ADP-ribose polymerase (PARP) inhibitors [9]. Genetic analyses of CRPCs have revealed that mutations in the HR genes *BReast Cancer type 1 (BRCA1;* 0.7%), *BReast CAncer type 2 (BRCA2;* 13.3%) and *ataxia telangiectasia mutated* (*ATM;* 7.3%) occur in a sizeable fraction of advanced CRPC patients [10]. This observation directly led to the 2020 approval of the PARP inhibitor Olaparib for advanced CRPC patients harboring a mutation in *BRCA1/2* or *ATM* [10, 11]. However, while Olaparib is approved in advanced *ATM* mutant CRPC, further studies and clinical data have demonstrated that this drug has marginal clinical efficacy on this patient cohort [12, 13]. The lack of utility of PARP inhibitors in *ATM* mutant prostate cancers is further reflected in the scope of the FDA approvals for two other PARP inhibitors, Rucaparib and Niraparib, both of which were approved for *BRCA1/2*-mutated metastatic CRPC but not *ATM* mutated patients [14–17]. Thus, there is a compelling need to identify alternative therapeutic avenues for *ATM* mutant prostate cancer patients, demanding a deeper understanding of the effects of ATM loss in prostate cancer.

Here, we sought to elucidate the impact of ATM loss on DDR in CRPC. Through unbiased phosphoproteomic analysis, we unveiled that ATM-deficient CRPC lines maintain a strong dependence on the ATM kinase pathway in response to DNA damage produced by ionizing radiation (IR), despite convincing loss of ATM kinase activity. We show that the kinases ataxia telangiectasia and Rad3-related (ATR), and DNA-dependent protein kinase catalytic subunit (DNA-PKcs) had significant redundancy in this pathway and hypothesized that these kinases were compensating for ATM-deficiency in this context. Indeed, dual inhibition of both kinases effectively attenuated DDR in response to IR and was able to impact replication fork dynamics in ATM-deficient CRPC lines. We leveraged this knowledge to employ the RUVBL1/2 ATPase inhibitor Compound B, which we previously demonstrated leads to the degradation of both ATR and DNA-PKcs [18]. In ATM-deficient CRPC cells, Compound B mimicked dual therapy in its ability to effectively radiosensitize and impact replication fork dynamics. Overall, our study suggests that Compound B may be a novel therapeutic strategy in ATM-mutant prostate cancers, particularly in combination with a DNA damaging agent such as IR.

## Results

### Generation and Validation of ATM-deficient CRPC lines

To characterize the mutational landscape of ATM in prostate cancer, we identified all available prostate cancer specimens in the cBioPortal database for putative driver mutations in ATM (n=206), excluding variants of unknown significance. We found that the majority (∼69.4%) of these mutations resulted in ATM protein truncation and were not localized to a particular hotspot, but rather distributed across the *ATM* locus as predominantly loss-of-function mutations **(Figure 1a)** [10, 19–36]. Notably, this finding was maintained independent of the dataset used **(Supplemental Figure S1)**. Thus, we decided knocking out ATM would best simulate a wide range of clinical scenarios. We employed CRISPR-Cas9 editing to generate clinically relevant paired ATM-proficient and ATM-deficient C4-2 and 22Rv1 CRPC cell lines (denoted V2 and KOs, respectively) and confirmed ATM protein loss in selected ATM-deficient lines via western blot **(Figure 1b)** [37]. Based on ATM’s known responses to DNA damage, we further confirmed loss of ATM kinase activity by observing loss of ATM Ser1981 autophosphorylation and an attenuation of the downstream ATM target pKAP1 (Ser824) in response to IR **(Figure 1c,d Supplemental Figure S2).**

**Figure 1:**
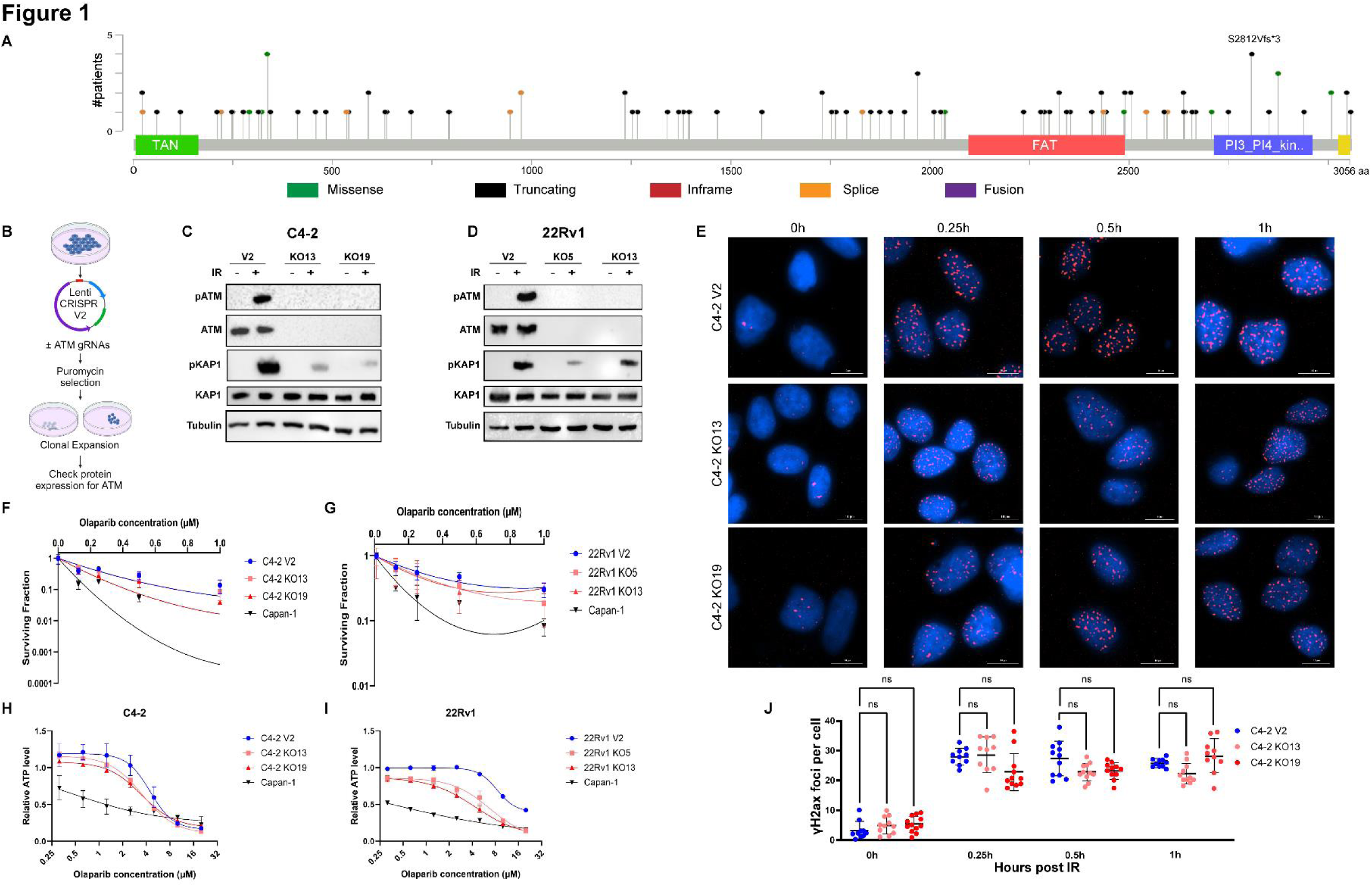
Generation and validation of ATM-deficient CRPC lines. **1a** Lollipop graph of known pathogenic mutation locations on the ATM gene in prostate cancer demonstrates that the majority of them are distributed throughout the gene and are often loss-of-function truncation mutations. Figure generated using data from cBioPortal on 1.15.2024; excluding variants of unknown significance. N=9424 samples from 9221 patients found 206 driver mutations in ATM. 143/206 (69.4%) of these are truncation mutations. Other types of mutations include missense (29/206; 14.1%), splice (21/206; 10.2%), and fusion mutations (13/206; 6.3%). See **Method**s section for specifics on study selection. **1b** Schematic of how C4-2 and 22Rv1 cell lines were generated via CRISPR-Cas9 using either the empty vector (LentiCRISPR.V2) plasmid, or LentiCRISPR.V2 with gRNA directed towards ATM (KOs). Two cell populations were selected per cell line. “V2” denotes the empty vector (ATM-proficient). For C4-2, KO13 and KO19 denotes the ATM-deficient clone 13 and 19, respectively. For 22Rv1, KO5 and KO13 denotes the ATM-deficient clone 5 and 13, respectively. Figure created on BioRender.com. **1c and 1d** Western blot shows knockdown of ATM protein expression, loss of ATM autophosphorylation (pSer1981) in response to DNA damage, and decreased downstream ATM kinase ability to phosphorylate KAP1 at Ser824 in response to IR in C4-2 **(1c)** and 22Rv1 **(1d)** prostate cell lines. Two clones were chosen per cell line based on efficiency of ATM loss on western blot. Each western was run on multiple gels with the same lysates. For ease of presentation, only one loading control is shown; ***Supplemental Figure S2*** shows the loading control for each independent gel. **1f and 1g** Clonogenic survival assays of both C4-2 V2 and ATM KO cells **(1e)** and 22Rv1 V2 and ATM KO cells **(1f)** demonstrates poor sensitivity of ATM KO lines to Olaparib as opposed to the *BRCA2*-mutant positive control line, Capan-1. Each experiment was done 3 independent times in technical triplicate; values presented represent mean ± SD for a selected independent experiment. Survival curves were generated using the linear quadratic equation (Y is fraction of cells surviving) on GraphPad Prism. **1h and 1i** CellTiter-Glo assays (9d) of both C4-2 V2 and ATM KO cells **(1g)** and 22Rv1 and ATM KO cells **(1h)** demonstrates poor sensitivity of ATM KO lines to Olaparib compared to Capan-1. Each experiment was done 3 independent times: two times in technical triplicate, and once in technical sextuplet. One independent experiment is presented. Values presented represent mean ± SD. Curve was generated running the non-linear [Inhibitor] vs. response—Variable slope (four parameters) dose response on GraphPad Prism. **1e and 1j** Time-course immunofluorescence (IF) staining of γH2AX foci recruitment in response to 1Gy IR on DMSO treated C4-2 V2, KO13, and KO19 cells demonstrates no statistically significant difference in number of γH2AX foci recruited up to 1 hr. Three independent experiments were run; data presented is from one experiment. Representative images were pre-processed for background flattening with consistent rolling ball diameter and subsequently deconvoluted using Gen5 software. Scale bar represents 10 µM **(1j)** Foci were quantified using the Spot Counting module on Imaris software. A minimum of 100 cells were quantified per condition in the selected experiment. Each dot represents an image field. Statistical significance was calculated with One-Way ANOVA with Tukey’s multiple comparison test.

Since clinical data have demonstrated that ATM mutant prostate cancers show limited sensitivity to PARP inhibitors, we next assessed the sensitivity of our ATM-deficient cell lines to the PARP inhibitor Olaparib with both clonogenic survival assays and CellTiter-Glo assays (**Figure 1f-i**) [12, 15, 16]. In each instance, we utilized the *BRCA2*-mutant pancreatic cancer cell line Capan-1 as a positive control, as it shows exquisite sensitivity to PARP inhibitors [38]. We found that both C4-2 and 22Rv1 ATM-deficient cells demonstrate only weak sensitization to Olaparib compared to ATM-proficient controls while Capan-1 cells were highly sensitive, which mirrors the limited clinical efficacy of PARP inhibitors in patients with ATM-mutated prostate cancer [12, 15, 17]. Since ATM is involved in initiating the DDR cascade, we interrogated whether ATM loss had an effect on γH2AX foci formation in response to 1 Gy IR. We found no significant difference in the kinetics of γH2AX foci formation in ATM-proficient and ATM-deficient cells at time points ranging from 15 min to 1 hr in both C4-2 and 22Rv1 cell lines (**Figure 1e,j**; Supplemental Figure S3), suggesting that ATM-deficient prostate cancer cells maintain the ability to initiate the DNA-damage response.

### Effective DNA damage repair in ATM-deficient prostate cancer depends on ATR and DNA-PKcs

To characterize how ATM-deficient prostate cancer cells maintain the ability to repair DNA double stranded breaks (DSBs) induced by IR, we performed an unbiased phosphoproteome enrichment coupled to quantitative mass spectrometry screen in response to irradiation. This approach can detect changes in DNA damage signaling that relies primarily on rapid protein phosphorylation upon the occurrence of DSBs [39]. We subjected C4-2 and 22Rv1 ATM-proficient and ATM-deficient cells to 10 Gy IR to induce DSBs, harvested the cells 1 hr later, and performed relative quantification using mass spectrometry-based phosphoproteomics for phosphopeptide identification – a timeline consistent with previously published studies on ATM in other model systems **(Figure 2a)** [40]. A total of 8,626 and 8,009 unique phosphopeptides were identified in C4-2 and 22Rv1 cells, respectively. Volcano plots comparing irradiated samples to non-irradiated controls in all lines demonstrated a robust response to IR with distinct panels of phosphopeptides **(Figure 2b,c,e,f; Supplemental Figure S4a).** Further, ATM-deficient cells exhibited differential responses to IR compared to ATM-proficient control cells **(Supplemental Figure S4b, S4c, S4d).**

**Figure 2:**
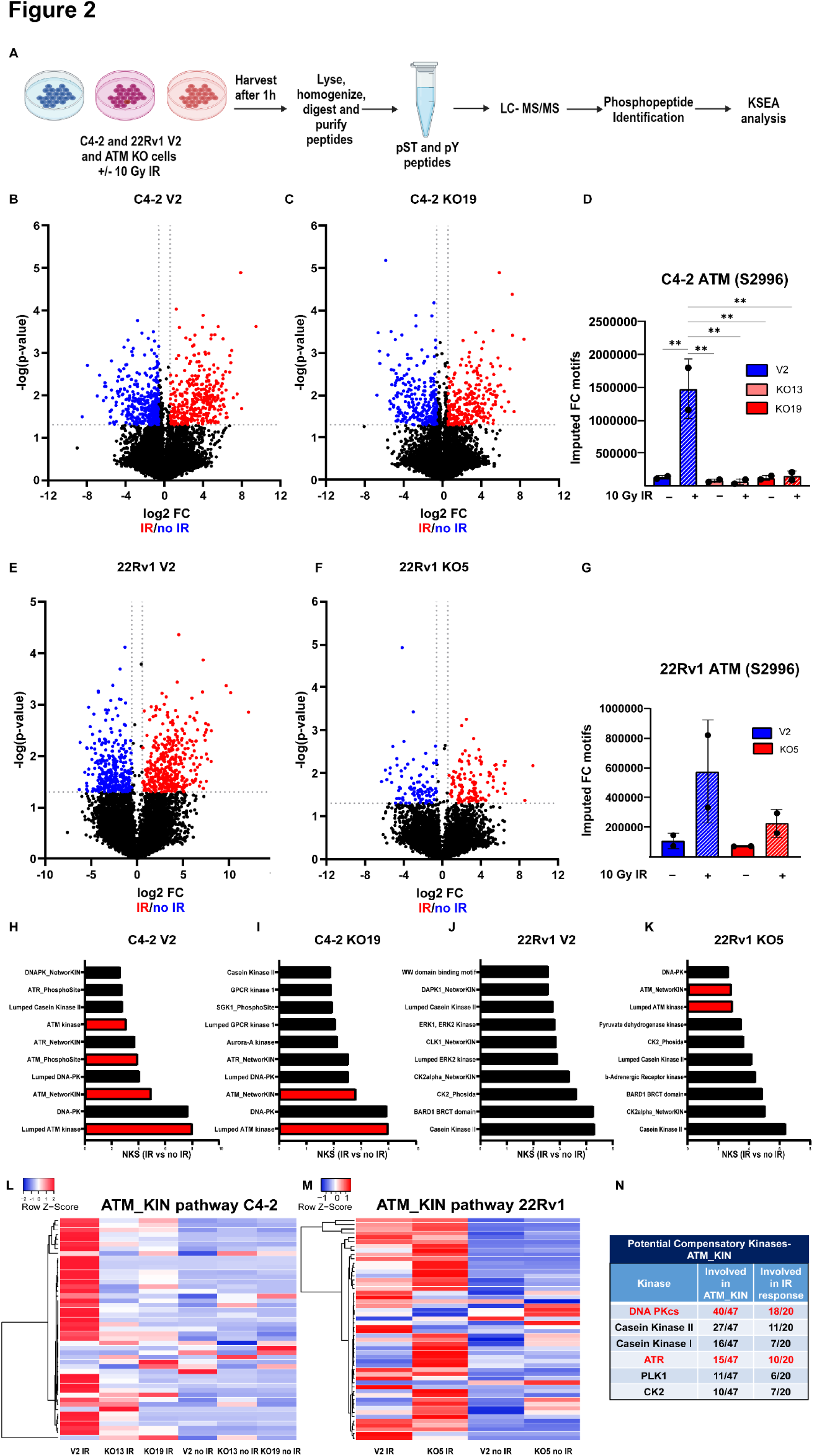
Effective DNA damage repair in ATM-deficient prostate cancer depends on ATR and DNA-PKcs. **2a** Schematic of unbiased phosphoproteomics. C4-2 and 22Rv1 V2 and KO cell lines were treated with ± 10 Gy IR and harvested 1 hr later. Samples were then processed for mass spectrometry and downstream analysis was performed via Kinase-Substrate Enrichment Analysis (KSEA). Figure adapted from Drake et. al, Cell 2016 [66] and created with BioRender.com. **2b and 2c** Volcano plots of C4-2 V2 **(2b)** and C4-2 KO19 **(2c)** demonstrate distinct panels of differentially expressed phosphosites in response to IR. Volcano plots were generated with Quantum-Normalized Log Transformed (QNLT) values. Vertical dotted line represents fold change cut off of 1.5. Horizontal dotted line represents p-value of 0.05. **2d** Analysis of pATM at Ser2996 demonstrates significant loss of ATM kinase activity in response to IR in C4-2 KO13 and KO19 cell lines. Note that pSer1981 was not available in the phosphoproteomic library for analysis. Statistical analysis generated with Tukey’s multiple comparison test (One-Way ANOVA). **2e and 2f** Volcano plots of 22Rv1 V2 **(2e)** and 22Rv1 KO5 **(2f)** demonstrate distinct panels of differentially expressed phosphosites in response to IR. Volcano plots were generated with Quantum-Normalized Log Transformed (QNLT) values. Vertical dotted line represents fold change cut off of 1.5. Horizontal dotted line represents p-value of 0.05. **2g** Analysis of pATM at Ser2996 demonstrates loss of ATM kinase activity in response to IR in the 22Rv1 KO5 cell line. Note that, as with C4-2 lines, pSer1981 was not available in the phosphoproteomic library for analysis. **2h-2k** Kinase Substrate Enrichment Analyses (KSEA) of C4-2 V2 **(2h),** C4-2 KO19 **(2i),** 22Rv1 V2 **(2j)**, and 22Rv1 KO5 **(2k)** cell lines, presented as irradiated samples/non-irradiated controls. The pathways were filtered for number of hits > 5 and FDR < 0.05. The top ten pathways, by Normalized Kolmogorov–Smirnov score (NKS score), are shown. Analyses demonstrate that despite convincing loss of ATM in ATM-deficient cell lines, there is still a dependence on the ATM pathway. Note that in 22Rv1 V2, the ATM_KIN pathway is still significantly upregulated but is not in the top 10 pathways. (***See Supplemental Figure S5***) **2l and 2m** Heatmaps of the ATM_KIN pathway in C4-2 **(2l)** and 22Rv1 **(2m)** shown as the average of n=2 samples used for phosphoproteomics. Heatmaps were generated with http://www.heatmapper.ca/expression/ using Average linkage clustering method and Euclidean distance measurement method. **2n** A table of the compensatory kinases involved in the ATM_kinase pathway and IR-dependent kinases identify DNA-PKcs and ATR as two of the top kinases. In the “Involved in ATM_KIN” column, the denominator corresponds to the total number of phosphopeptides involved in the ATM_KIN pathway. The numerator corresponds to the number of phosphopeptides that overlap between the chosen pathway and the ATM_KIN pathway. In the “Involved in IR response” column, the denominator refers to the total number of phosphopeptides in the ATM_KIN pathway that are important in the response to IR (as defined as a FC >2 with IR in KO19 C4-2 cells). The numerator corresponds to the number of phosphopeptides that overlap between the chosen pathway and the ATM_KIN pathway in response to IR.

Kinase-Substrate Enrichment Analysis (KSEA) confirmed that phosphopeptide substrates involved in the ATM kinase pathway were enriched in C4-2 and 22Rv1 ATM-proficient controls compared to matched ATM-deficient cells in response to IR (Figure S4f-h). Further, in ATM-deficient cells, ATM autophosphorylation on the DNA-damage inducible site Ser2996 was abrogated, confirming loss of ATM kinase activity in these models (**Figure 2d,g**) [41].

Despite convincing loss of ATM kinase expression and activity, KSEA demonstrated that the ATM-deficient cell lines were still enriched for ATM kinase pathway phosphopeptide substrates in response to IR, i.e. ATM-deficient cells maintained dependence on the downstream targets of the ATM kinase signaling cascade (**Figure 2h-k**, Supplemental Figure S4e, S5). Heatmap analysis of the ATM_KIN pathway further supported this finding. Phosphopeptide substrates involved in this pathway were modestly increased in irradiated ATM-deficient C4-2 cells compared to non-irradiated controls, and irradiated 22Rv1 ATM-deficient cells remained comparable to ATM-proficient cells in response to IR (**Figure 2l,m**). Thus, these findings suggest functional redundancy in the DDR signaling cascade likely via compensatory kinases that maintained phosphorylation of ATM kinase pathway targets.

To elucidate which kinases might be potentially responsible for this compensatory effect, we identified all kinases that were known or predicted to phosphorylate phosphopeptides involved in the ATM_KIN pathway. We further filtered phosphopeptides in the ATM_KIN pathway upregulated in ATM-deficient cells in response to IR (as defined by phosphosites with a fold change ≥ 2 in response to IR in C4-2 KO19 cells) (**Figure 2n**) This analysis identified ATR and DNA-PKcs as top potential compensatory kinases, which both have known functions in DNA damage repair and have targeted inhibitors in ongoing clinical trials [42–45].

### Dual inhibition of ATR and DNA-PKcs impairs DNA-Damage repair in ATM-deficient prostate cancer cells

To assess whether ATM-deficient prostate cancer cell lines rely on ATR or DNA-PKcs for efficient repair of DSBs, we performed clonogenic survival assays in response to IR. We pretreated the cells with either the selective ATR kinase inhibitor VX-970, the selective DNA-PKcs inhibitor M3814, or their combination, and subjected them to escalating doses of IR [45–47]. We observed minimal attenuation of colony formation with VX-970 and only a modest effect with M3814 compared to DMSO control in ATM-deficient cells in response to IR. Notably, dual inhibition of ATR and DNA-PKcs markedly radiosensitized ATM-deficient cells **(Figure 3a,b; Supplemental Figure S6a-c)**. We next confirmed that the observed radiosensitization could be replicated with alternative inhibitors; replacing VX-970 with the highly selective ATR inhibitor, BAY1895344 or substituting M3814 with the selective DNA-PKcs inhibitor, NU7441 yielded similar results **(Supplemental Figure S6d-h)** [48, 49]. Next, to evaluate the ability of ATM-deficient cells to initiate the DNA damage signaling cascade, we assessed the impact of ATR or DNA-PKcs inhibition on γH2AX foci formation in response to 1 Gy IR. We found that VX-970 or M3814 treatment alone was insufficient to reduce γH2AX foci formation 1 hr after 1 Gy IR in ATM-deficient cells. However, dual inhibition significantly decreased the number of γH2AX foci in the ATM-deficient cell lines **(Figure 3c,d; Supplemental Figure S7)**. Analysis of the kinetics of γH2AX foci demonstrated that this was due to reduced formation of γH2AX by 1 hr rather than a product of foci resolution **(Supplemental Figure S8)**.

**Figure 3:**
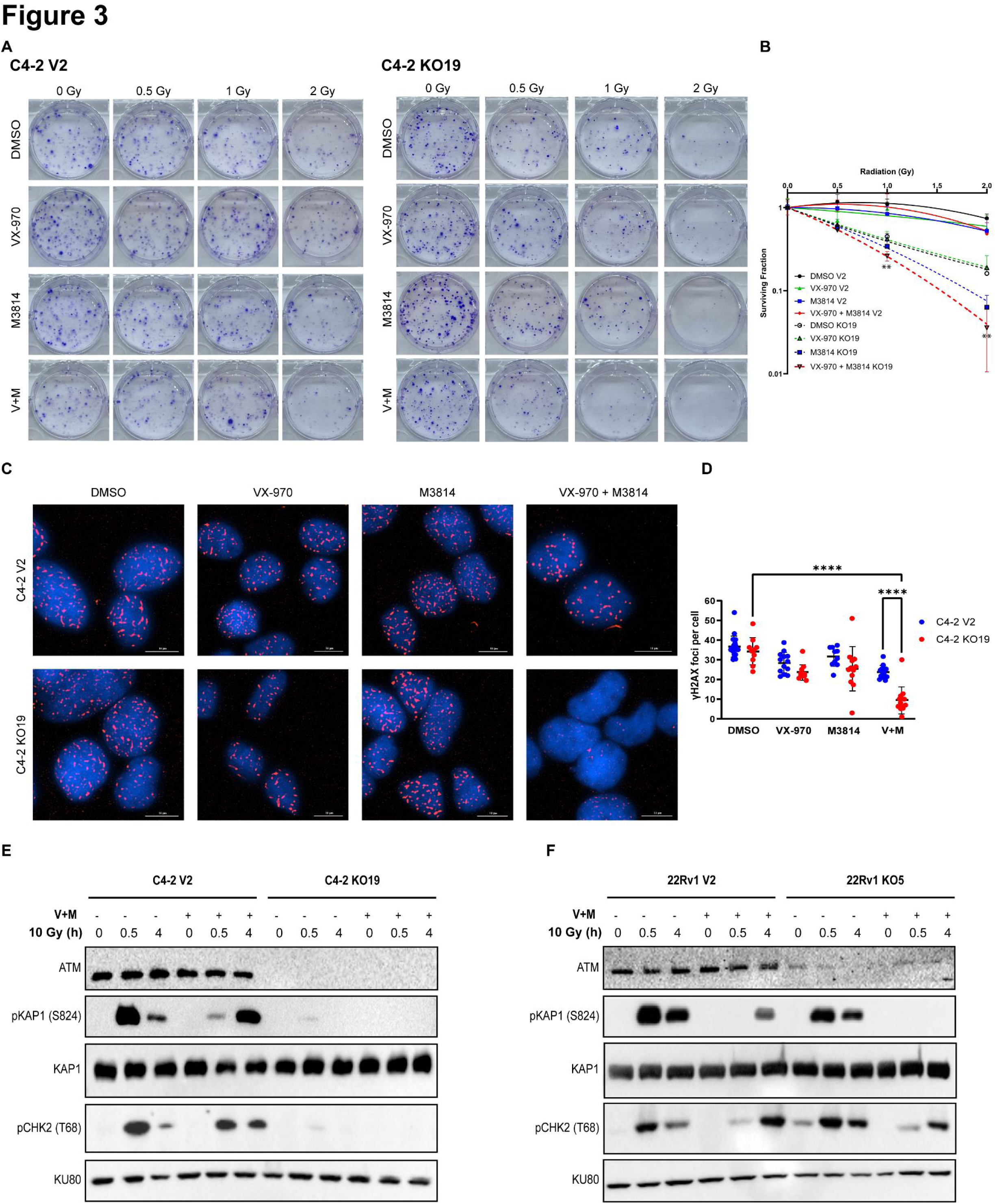
Dual inhibition of ATR and DNA-PKcs is required to impair DNA-Damage repair in ATM-deficient prostate cancer cells. **3a and b** Clonogenic survival assay of C4-2 V2 and C4-2 KO19 with 200 nM M3814 and 10 nM VX-970 and escalating doses of irradiation demonstrates that combination therapy is most effective in the context of ATM loss **(3a)**. Quantified values **(3b)** represent mean ± SD in technical triplicate. Colonies were quantified with ImageJ cell counter plugin, and survival curve was generated using the linear quadratic equation (Y is fraction of cells surviving) on GraphPad Prism. Statistics shown analyze C4-2 KO19 DMSO vs. C4-2 KO19 VX-970 + M3814; determined by multiple t-tests with the Holm-Šídák method. **3c** IF staining of C4-2 V2 and C4-2 KO19 cells for γH2AX foci. Cells were pre-treated with DMSO control, 1 µM VX-970, 10 µM M3814, or 1 µM VX-970+10 µM M3814 for 1.5 hrs, subjected to 1 Gy IR, and harvested 1 hr later. γH2AX foci are not observed at 1 hr in the combination therapy group. Scale bar represents 10 µm. Three independent experiments were run; data presented is from one experiment. Representative images were pre-processed for background flattening with consistent rolling ball diameter and subsequently deconvoluted using Gen5 software. **3d** γH2AX foci were quantified using the Spot Counting module on Imaris software. A minimum of 70 cells were quantified per condition in the selected experiment. Each dot represents an image field. Error bars represent mean ±SD. Quantification of γH2AX foci per cell demonstrates a statistically significant reduction in combination therapy in KO19 as opposed to DMSO control. An Ordinary One Way ANOVA with Turkey’s multiple comparison test was used for statistical analysis. (**** represents p-value <0.0001). **3e** Western blot of C4-2 V2 and KO19 cells demonstrates that combination therapy attenuates residual downstream ATM kinase activity in ATM-deficient cells. Cells were treated with either DMSO or a combination of 1 µM VX-970 and 10 µM M3814 for 1.5 hrs, subsequently subjected to 10 Gy IR, and harvested at the indicated time points. 0h time point represents no IR. Western was run with the same lysates on multiple gels, ***Supplemental Figure S9a*** shows loading controls for each gel. In C4-2 KO19 cells, the residual phosphorylation of downstream ATM targets pKAP1 on Ser824 and pCHK2 on Thr68 is greatly decreased with dual therapy. C4-2 V2 cells demonstrate delayed kinetics of KAP1 and CHK2 phosphorylation. **3f** Western blot of 22Rv1 V2 and KO5 cells demonstrates that combination therapy attenuates residual downstream ATM kinase activity in ATM deficient cells. Experiment is as described in **3e**. Western was run with the same lysates on multiple gels, ***Supplemental Figure S9b*** shows loading controls for each gel. In 22Rv1 KO5 cells, the residual phosphorylation of downstream ATM targets pKAP1 (Ser824) is abolished with dual therapy and pCHK2 (Thr68) is delayed and decreased.

We next interrogated if dual inhibition of ATR and DNA-PKcs could prevent residual downstream signaling of the ATM pathway. We pre-treated ATM-proficient and ATM-deficient cells with either DMSO or a combination of VX-970 and M3814 for 1.5 hrs, subjected them to 10 Gy IR, and harvested them 0.5 hrs and 4 hrs after irradiation. In C4-2 cells, we noted that dual inhibition abrogated residual phosphorylation of the downstream ATM pathway targets KAP1 (Ser824) and CHK2 (Thr68), whereas there was only a moderate effect in C4-2 ATM-proficient controls **(Figure 3e, Supplemental S9a)**. In 22Rv1 ATM-deficient cells, we observed a complete loss of pKAP1 and a substantial delay and attenuation of pCHK2 with dual inhibition. We also found that dual kinase inhibition in ATM-proficient 22Rv1 cells impaired ATM downstream signaling, albeit to a weaker extent than observed for ATM-deficient 22Rv1 cells **(Figure 3f, Supplemental S9b)**. This finding mirrored our observations for the effects of dual kinase inhibition on γH2AX foci formation, with dual kinase inhibition providing the greatest effect **(Figure 3c,d, Supplemental Figure S7)**.

### Dual inhibition of ATR and DNA-PKcs affects replication fork dynamics

With the knowledge that DNA breaks are not limited to DSBs created by IR, and that any unrepaired breaks might affect the fidelity of DNA replication, we investigated the repercussion of single or dual inhibition of ATR and DNA-PKcs on replication fork dynamics in ATM-deficient C4-2 KO19 cells [50, 51]. To assess these effects, we employed DNA fiber assays: cells were pre-treated with DMSO, VX-970, M3814, or both inhibitors for 1.5 hrs, followed by sequential pulse-labeling with two thymidine analogs, iodo-deoxyuridine (IdU; in red) and chloro-deoxyuridine (CldU; in green) for 20 min each. Dual inhibition of ATR and DNA-PKcs led to reductions in replication fork length and fork speed compared to vehicle or single inhibitor treatment, while maintaining replication fork symmetry **(Figure 4; Supplemental Figure S10a)**. These findings suggest that dual ATR and DNA-PKcs inhibition affected replication fork progression.

**Figure 4:**
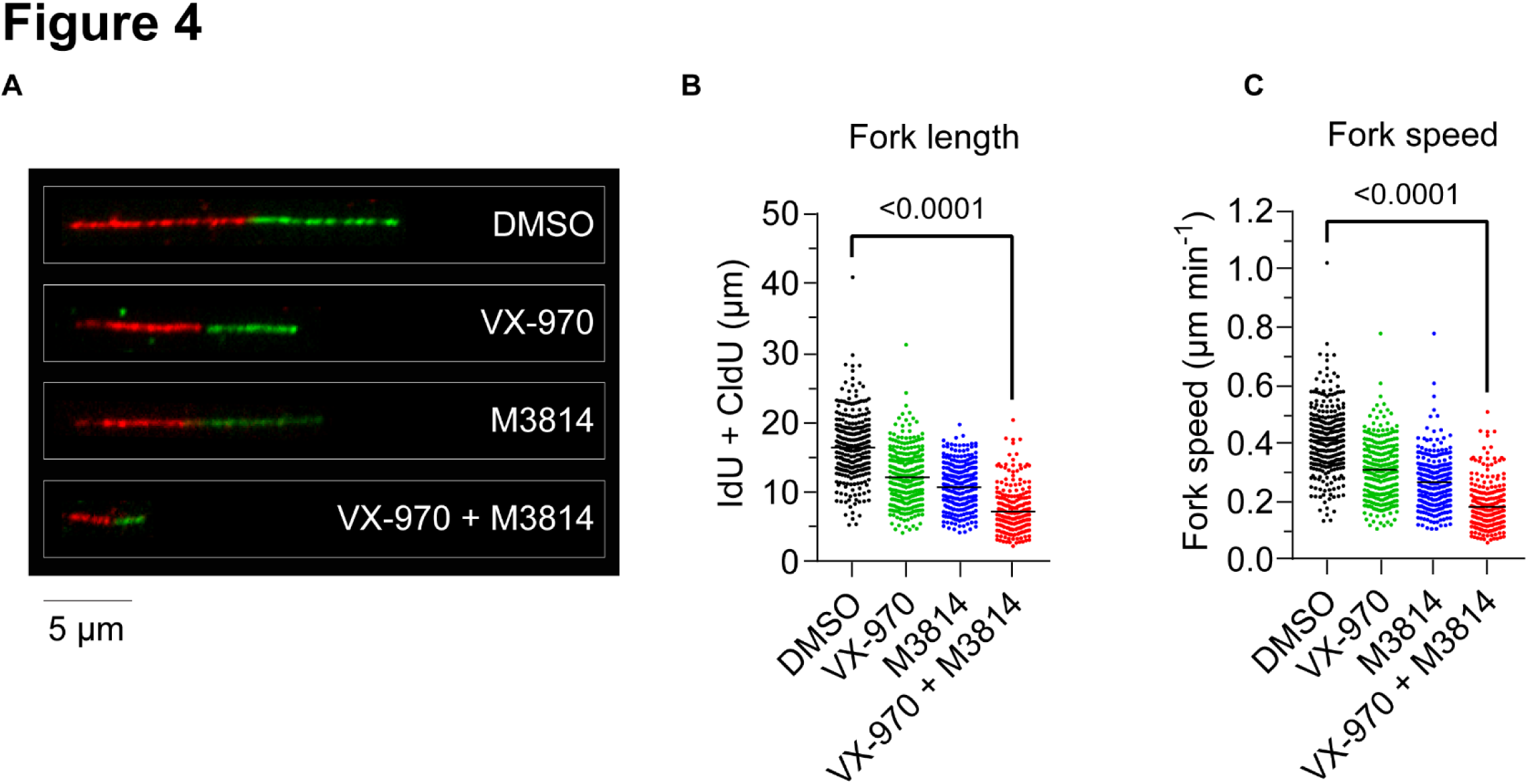
Dual inhibition of ATR and DNA-PKcs affects replication fork dynamics. **4a** Representative fibers from fiber assay. C4-2 KO19 cells were plated and treated 48 hrs later with DMSO, 1 µM VX-970, 10 µM M3814, or a combination for 1.5 hours. Cells were pulse labeled with IdU (red) and CIdU (green) for 20 min each, lysed and stretched on a slide, and imaged. Scale bar represents 5 µm. **4b and c** Quantification of fiber fork length and fork speed. Data are mean ± S.E.M. from DMSO, n=300; VX-970, n= 300, M3814, n= 301; VX-970 + M3814, n=234 fibers pooled from 2 independent experiments. Statistics calculated by Kruskal-Wallis multiple comparison test.

### RUVBL1/2 inhibition as a novel therapeutic avenue for ATM-deficient prostate cancer

While our findings demonstrate the need for dual kinase inhibition in ATM-deficient prostate cells, these kinase inhibitors have been shown to have significant adverse events, and we expect the combination to only exacerbate these symptoms [42, 43, 45]. To circumvent the potential toxicity of utilizing two kinase inhibitors, we next sought to determine if the previously published RUVBL1/2 ATPase inhibitor, Compound B, could be utilized in ATM-deficient prostate cancer cells. We have previously demonstrated in lung cancer models that Compound B inhibits the ATPase activity of RUVBL1/2, leading to inhibition of PAQosome maturation. The PAQosome is required for stability of the phosphatidylinositol 3-kinase-related kinases (PIKK) family of proteins, including ATR and DNA-PKcs. Thus, Compound B treatment leads to downstream depletion of ATR and DNA-PKcs protein levels. Importantly, our group previously demonstrated that this was cancer selective, as normal lung epithelial cells were spared [18, 52]. To confirm that Compound B treatment maintained the ability to deplete the levels of ATR and DNA-PKcs in C4-2 and 22Rv1 ATM-proficient and ATM-deficient lines, we pretreated the cells with 100 nM Compound B or its inactive enantiomer Compound C for 72 hours (a timeline we previously established for efficient PIKK depletion) and assessed protein levels by western blot. We confirmed that Compound B treatment depleted ATR and DNA-PKcs kinase protein levels in all cell lines **(Figure 5a,b)**. Next, we tested if RUVBL1/2 inhibition affected the ability of prostate cancer cell lines to effectively repair DSBs via clonogenic survival assays in response to IR. Pretreatment with Compound B followed by exposure to escalating doses of IR markedly radiosensitized both ATM-proficient and ATM-deficient cells compared to pretreatment with the control Compound C **(Figure 5c,d,h,i)**.

**Figure 5:**
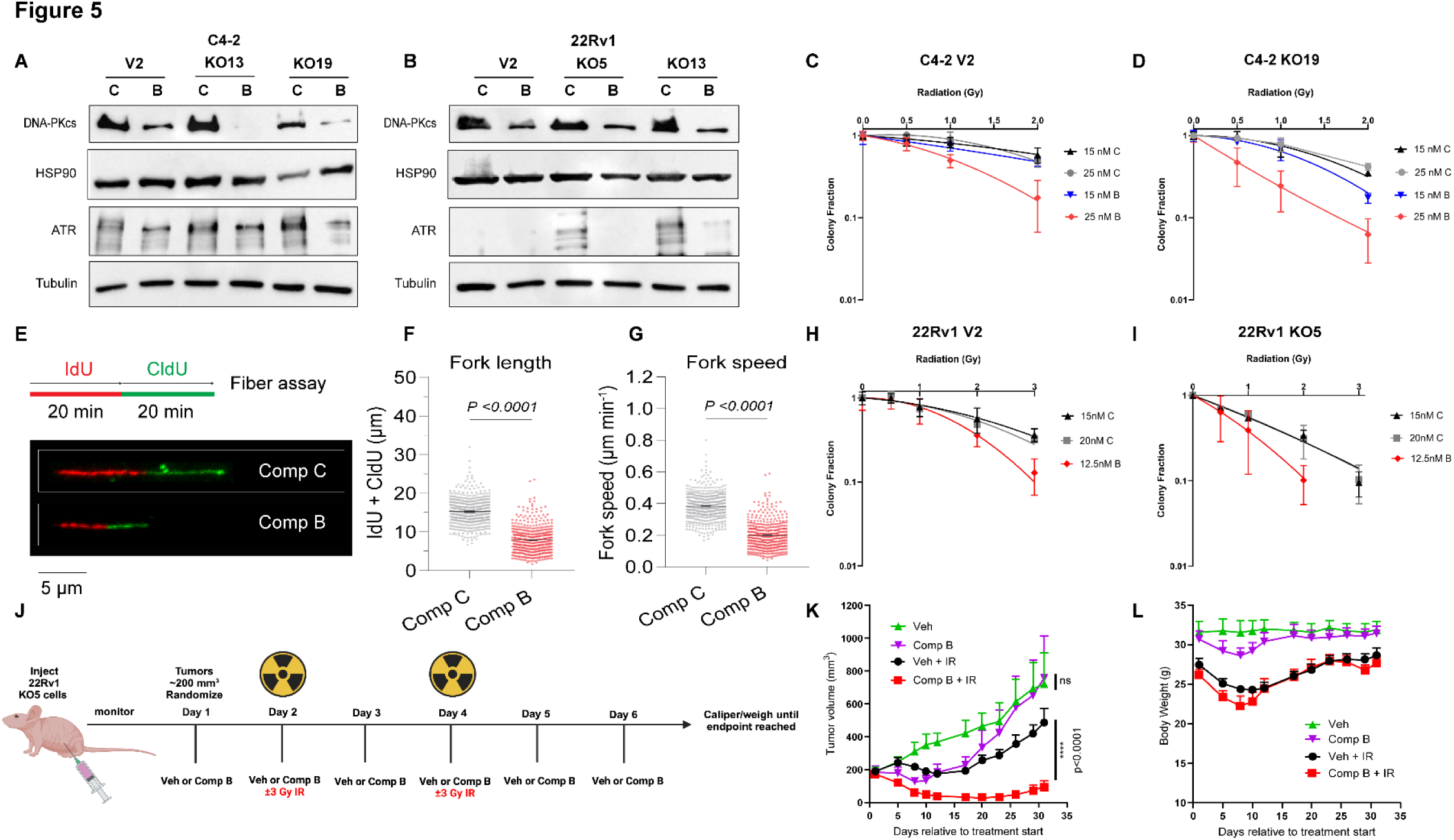
Compound B as a novel therapeutic in ATM-deficient prostate cancer. **5a and b** Western of C4-2 **(5a)** and 22Rv1 **(5b)** in ATM-proficient and deficient cells demonstrates that Compound B treatment successfully depletes protein levels of ATR and DNA-PKcs. Cells were pre-treated with either 100 nM Compound B or 100 nM Compound C (control enantiomer) 72 hrs before harvesting to ensure PIKK depletion. **5c-d, 5h-i** Clonogenic survival assays of C4-2 V2 **(5c)**, C4-2 ATM KO19 **(5d)**, 22Rv1 V2 **(5h),** and 22Rv1 KO5 **(5i)** demonstrates the ability of Compound B to radiosensitize *in vitro*. 2-3 biological replicates, each in technical triplicate were performed per cell line; data presented is from one independent experiment in technical triplicate. Data presented as mean ± SD. Colonies were quantified with ImageJ cell counter plugin, and survival curve was generated using the linear quadratic equation (Y is fraction of cells surviving) on GraphPad Prism. **5e-g** Fiber assays performed on C4-2 KO19 cells. Cells were treated with 100 nM Compound B or Compound C for 72 hours. Representative fibers are shown in **5e.** A decrease in fork length **(5f)** and fork speed **(5g)** was noted with Compound B treatment. Data are mean ± S.E.M from n=450 (Compound B) and n=451 (Compound C) forks pooled from 3 independent experiments. Statistics were calculated with Mann-Whitney U test (unpaired and nonparametric). **5j-l** *In vivo* animal experiments demonstrate the ability of Compound B to radiosensitize. 22Rv1 KO5 cells were xenografted into nude mice and when tumor volume reached ∼200 mm^3^ mice were randomized. Schematic of treatment outlined in **5j,** created with BioRender.com. Raw tumor volumes **(5k)** and mouse weight **(5l)** were monitored, and are presented as mean ± S.E.M. Statistical significance between vehicle and Compound B, and between vehicle+ IR and Compound B+ IR were assessed via 2-Way ANOVA with Bonferroni’s multiple comparison’s test. N=5 mice were included in each group.

To determine if Compound B influenced replication fork progression in ATM-deficient prostate cancer cells, we performed DNA fiber assays following pretreatment of C4-2 KO19 cells with Compound B or C for 72 hours. In agreement with dual inhibition with VX-970 and M3814 **(Figure 4)**, fork length and fork speed were significantly decreased and fork symmetry was modestly affected with Compound B treatment, indicating Compound B was able to affect replication fork progression **(Figure 5e-g; Supplemental Figure S10b)**. Importantly, treatment with Compound B decreased fork length to the same extent as dual inhibition of ATR and DNA-PKcs with VX-970 and M3814 **(Supplemental Figure S10c)**.

We next assessed the therapeutic potential of Compound B *in vivo* using IR to induce DNA damage. We xenografted 22Rv1 V2 or KO5 cells into the right flank of male nude mice. When the tumor volume reached ∼200mm^3^, we randomized the animals to receive vehicle (PEG200), 125 mg/kg Compound B, or Vehicle/Compound B with IR. Compound B treatment was given daily (in two doses of 62.5 mg/kg each) for 6 days, with ± 3 Gy IR directly targeted to the tumor on days 2 and 4. This dosing schedule allowed Compound B to initiate PIKK depletion before the first dose of IR, and allowed ATR and DNA-PKcs inhibition to persist for days after generation of DSBs with IR **(Figure 5j)**. We found that Compound B in combination with IR significantly attenuated the growth of ATM-deficient 22Rv1 KO5 cells **(Figure 5k)**. While the combination of Compound B and IR initially caused moderate weight loss suggest, the mice rapidly regained weight **(Figure 5l)**. In ATM-proficient 22Rv1 V2 xenografts, although combination of Compound B was most effective, the effect was modest and no significant difference was seen between Compound B+ IR and IR alone. **(Supplemental Figure S11)**. We note that there is a limited therapeutic window with this combination, as toxicity can be observed with higher doses. Taken together, these data suggest Compound B could be a novel therapeutic strategy in ATM-deficient prostate cancer in combination with IR.

## Discussion

Genomic analyses of advanced prostate cancers revealed that a significant percentage of patients harbor loss-of-function mutations in *ATM* [10, 20, 53]. While preliminary data initially supported the use of PARP inhibitors in this context, clinical data has demonstrated limited efficacy in this patient population, necessitating the need for alternative therapeutic modalities [12, 22]. Here, we interrogated how ATM loss impacted the ability of CRPC cell line models to respond to and effectively repair irradiation-induced DNA damage to identify targetable vulnerabilities in these cancers.

We successfully generated ATM-deficient models that demonstrated limited sensitivity to the PARP inhibitor Olaparib. This finding mirrors clinical studies that have demonstrated that BRCA2 loss confers exquisite sensitivity to PARP inhibitors, while ATM loss only confers limited sensitization [12, 13, 38]. Further, we found that for ATM-deficient CRPC cell lines γH2AX foci formation was not significantly impaired in response to DNA damage via IR. Thus, our ATM-deficient CRPC models accurately imitate clinical scenarios of ATM loss, while still maintaining their ability to effectively repair damaged DNA. Through unbiased phosphoproteomic analyses in our matched ATM-proficient and ATM-deficient prostate cells, we found that ATM-deficient cells maintain ATM kinase downstream signaling in response to DNA-damage by IR, despite the loss of ATM-kinase activity. When analyzing potential kinases that could complement for ATM loss via phosphorylation of its downstream targets, ATR and DNA-PKcs emerged as two top potential compensatory kinases. We focused on ATR and DNA-PKcs based on their extensively characterized association with ATM kinase in other cell models, in which many studies have pointed to considerable crosstalk that occurs between these three kinases [39, 54–58]. Further, our findings in CRPC cells are consistent with a phosphoproteomic study, in which lymphoblastoid ATM-deficient cell lines were shown to heavily depend on ATR and DNA-PKcs kinases when DNA breaks were induced with the radiomimetic drug, neocarzinostatin [59].

In our ATM-deficient models, we noted that dual inhibition of ATR and DNA-PKcs was most effective in preventing effective DNA damage repair. We observed that dual inhibition attenuated colony formation in response to IR to a greater extent than single agent treatment and was able to significantly impair γH2AX foci formation. Dual inhibition also optimally attenuated residual phosphorylation of downstream ATM targets pKAP1 and pCHK2. These findings indicate that the activity of any one of the two kinases is sufficient to mediate DDR in ATM-deficient cells, and that blockade of both ATR and DNA-PKcs is needed to prevent effective DDR in this context. Further, DNA fiber assays demonstrated a significant reduction in fork length and fork speed with dual inhibition compared to control or single inhibition, suggesting that dual therapy impacts replication fork dynamics. These findings suggest a dual pronged approach for combination therapy, in the impairment of DDR in response to DNA damage and in affecting DNA replication.

While our results indicate a requirement for dual targeting of ATR and DNA-PKcs in ATM-deficient prostate cancers, there are significant concerns about the clinical utility of combining two kinase inhibitors due to potential toxicity. In a Phase I study on solid tumor patients with the ATR inhibitor VX-970, 88.2% of patients experienced an adverse event (AE), and in combination with a DNA damaging agent, the AEs increased to 91.3% [43]. Studies with the newer, highly selective ATR inhibitor, BAY1895344, demonstrated significant hematologic toxicities, including anemia, neutropenia, and thrombocytopenias [42]. Further, in Phase I clinical trials of the DNA-PKcs inhibitor M3814, 71% of patients had at least one AE with treatment, with the most common toxicities being nausea, fatigue, vomiting, and pyrexia. Of these patients, 23% had Grade 3 adverse events, with maculopapular rash being common [44]. We would expect to exacerbate these events with the combination of the two therapies, which would significantly limit the clinical utility of this modality. To overcome this limitation, we utilized the RUVBL1/2 inhibitor, Compound B, which has been previously published to effectively deplete ATR and DNA-PKcs kinases in lung cancer models [18]. We were able to recapitulate loss of ATR and DNA-PKcs protein expression in our prostate models, and confirmed that Compound B was able to radiosensitize. Further, we confirmed that Compound B impacted replication fork dynamics to the same extent as dual kinase inhibition.

To assess the therapeutic potential of Compound B, we assessed its effectiveness *in vivo*. We noted that, in combination with IR, Compound B was able to significantly reduce tumor growth in 22Rv1 ATM-deficient xenograft models. We noted a greater radiosensitization effect in ATM-deficient cells compared to the ATM-proficient 22Rv1 V2 cell line. This indicates that Compound B could be utilized as a novel therapy in ATM-deficient prostate cancers, especially in combination with a DNA-damaging agent such as IR, and that Compound B might have greater efficacy in the ATM-deficient context. We have previously demonstrated that the resulting PIKK depletion from Compound B is cancer-selective and spares normal epithelial cells [18]. Thus, the radiosensitizing effects that we found are likely tumor-intrinsic, decreasing the potential for systemic toxicities associated with dual kinase inhibition.

## Materials and Methods

### Cell lines and culture conditions

C4-2 and 22Rv1 cells were obtained from American Type Culture Collection (ATCC). Their generated derivatives (22Rv1 and C4-2 V2 and ATM KO cells) were cultured in Roswell Park Memorial Institute (RPMI) 1640 medium (Sigma) supplemented with 10% Fetal Bovine Serum (FBS) (R&D Systems) 1% penicillin/streptomycin (P/S) (Sigma) and 0.1% normocin (Fisher Scientific) and maintained in 5 µg/mL puromycin (Fisher Scientific). Capan-1 cells were a kind gift from Rolf Brekken (UTSW) and maintained in RPMI 1640 with 10% FBS, 1% P/S, and 0.1% normocin. All cells were cultured in in 37 °C, 5% CO2. Cell lines were authenticated using short-tandem repeat (STR) profiling in the UTSW sequencing core and were periodically tested for mycoplasma via either MycoAlert (Lonza), MycoStrip Mycoplasma Detection kit (InvivoGen), or PlasmoTest Mycoplasma Detection Kit (InvivoGen).

### Inhibitors/Drugs Used

Olaparib (PARP inhibitor) was purchased from MedChemExpress (HY-10162). M3814 (DNA-PKcs inhibitor), VX-970 (ATR inhibitor), BAY1895344 (ATR inhibitor), and NU7441 (DNA-PKcs inhibitor) were purchased from Sellekchem (cat nos. S8586, S7102, S8666, and S2638 respectively). Puromycin was purchased from Fisher Scientific (Gibco, cat #A1113803). Compound B (RUVBL1/2 inhibitor) and Compound C (Compound B control) were synthesized by Daiichi-Sankyo.

### cBioPortal data

Data was accessed on 1.15.24. For Figure 1a, all datasets in the cBioPortal database were utilized, except for the database specific to Neuroendocrine Prostate Cancer. Additionally, only the TCGA (Firehouse) dataset was used of the TCGA datasets to avoid patient overlap between datasets. To generate the lollipop graphs, only driver mutations were plotted and variants of unknown significance were excluded. Both somatic and germline mutations were plotted. A total of 9424 samples corresponding to 9221 patients are included in Figure 1a.

### Transformation and Viral Infection

Competent Stbl3 bacteria were transformed with the LentiCRISPR.V2 vector with or without target sgRNA for ATM according to the LentiCRISPR.V2 cloning protocol from the Zhang lab [37]. Virus was produced in 293T cells. C4-2 and 22Rv1 cells were treated with virus and 1:1000 polybrene for 24hrs. Transfected cells were subsequently selected with 5 µg/mL puromycin. Cells were clonally selected twice and selected based on criteria of the most convincing ATM knockout on western blot after two rounds of selection. While an attempt was made for a single clone, the 22Rv1 cells are a somewhat mixed population and it must be noted that there is a small amount of ATM contamination.

Previously published CRISPR-Cas9 sgRNA sequences were utilized [60]:

ATM gRNA1_1 5’-CACCGCCAAGGCTATTCAGTGTGCG-3’

ATM gRNA1_2 3’-CGGTTCCGATAAGTCACACGC CAAA -5’

### CellTiter-Glo Assays

For CellTiter-Glo (CTG) experiments, 2.0-4.5 x 10^4^ cells per well were plated in a 96 well plate 1 day before treatment. The cells were treated with varying concentrations of Olaparib for 9 days and relative viability was assessed according to the CellTiter-Glo 2.0 (Promega) protocol. All experiments were done in technical triplicate or sextuplet and biological triplicate. Curve was generated via the non-linear [Inhibitor] vs. response-Variable slope (four parameters) dose response on GraphPad Prism (v10.1.2).

### Clonogenic Survival Assays

For clonogenic survival assays, 500-5,000 cells were plated in triplicate for each condition in 6-well plates. *For experiments performed with olaparib:* Plates were treated with the varying doses of Olaparib when cells adhered to the plate and harvested 14-18 days later. *For experiments performed with irradiation and ATR and DNA-PKcs inhibitors:* Cells were treated with indicated concentrations of inhibitors and were subsequently treated with varying doses of IR ∼1.5 hrs later. Plates were harvested 14-18 days later. *For experiments performed with irradiation and Compound B/C:* Cells were treated with either Compound B or Compound C in varying doses. Three days later, cells were subjected to varying doses of IR.

To harvest, cells were stained with crystal violet in 10% formalin and subsequently washed with water. Plates were allowed to dry and pictures were acquired. Quantification was performed using the ImageJ cell counter plugin. Survival curves were generated using the linear quadratic equation (Y is fraction of cells surviving) on GraphPad Prism (v10.1.2).

### Western Blotting

For experiments with IR alone, cells were treated with 10 Gy IR and subsequently harvested 1 hr later. For experiments with VX-970 and M3814, cells were pre-treated with 1 μM VX-970, 10 μM M3814, a combination, or DMSO control for 1.5 hrs, subsequently irradiated with 10 Gy IR, and harvested at the time point indicated. The 0h time point is not irradiated. For experiments with Compound B/C, cells were treated with 100 nM Compound B or 100 nM Compound C and harvested 72 hrs after to allow for PIKK depletion. **Harvesting:** Cells were scraped in PBS on ice and then spun down at 1500 rpm for 5 min. Cell pellet was resuspended in RIPA buffer (ThermoFisher Scientific) with protease (ThermoFisher Scientific) and phosphatase inhibitors (Millipore Sigma) and allowed to lyse on ice for 30 min with 30 sec vortex every 10 min. Lysates were then centrifuged at 14,000 rpm and protein-containing supernatant was taken as sample. The relative protein concentration was quantified using BCA (ThermoFisher Scientific) and mixed with 4X Lamelli buffer (BioRad). **Gel:** Lysates were boiled for 3-5 min immediately prior to loading the gel, and ∼40 µg of protein was loaded per gel. Homemade 4% stacking and 6% running gels were run at 60-80V in Tris/Glycine/SDS buffer (BioRad). Transfer was done overnight in 10% methanol in Tris/Glycine buffer (BioRad) onto either PVDF or Nitrocellulose membranes. Gels were blocked in 5% milk in TBS-T and primary antibodies (see Key Resources table for specifications) were incubated O/N on a rotator at 4°C. Membrane was washed 3X with TBS-T. Secondary was performed with either anti-mouse IgG (Cell Signaling Technology) or anti-rabbit (Cell Signaling Technology) IgG HRP-linked secondary antibody at 1:2000 dilution in 5% milk for an hour at RT. Membrane was washed 3X with TBS-T and then imaged.

### Phosphoproteomic Analysis

C4-2 and 22Rv1 V2 and KO cells were subjected to 10 Gy IR. Cells were harvested 1 hr post IR in PBS containing Mg^2+^ and Ca^2+^, with protease and phosphatase inhibitors. One sample was treated with sodium-orthovanadate (MedChemExpress) 15 min prior to harvest to establish a baseline for tyrosine phosphorylation. **Sample Preparation and Protein Extraction from cell pellets:** Cell pellets were lysed using buffer containing 7M urea, 2M thiourea, 0.4M Tris pH 8.0, 20% acetonitrile (ACN), 10 mM TCEP, 25 mM chloroacetamide, Thermo Scientific’s Halt protease inhibitor cocktail 1x concentration (originally 100x), and phosphatase inhibitors (HALT from Thermo). After adding 500 µL of the lysis buffer to the cell pellet, samples were placed on ice, then vortexed and centrifuged at 12,000 x g for 10 min. The samples were then sonicated for 5 seconds using a probe sonicator set at 30% amplitude and kept on ice during the entire sonication process. After sonication, the samples were incubated for 0.5 hrs at 37°C, then at room temperature for 15 min to reduce and alkylate cysteines and centrifuged at 12,000 x g for 10 min at 18°C. Protein concentration was measured using Bradford Assay (Bio-Rad). We then used 2.5 mg of protein, added 10 μg of 20 μg/mL Lysyl Endopeptidase (WAKO, 125-05061) and incubated the samples @ room temperature for 5-6 hr at pH 7.4. Then adjusted pH to 7.5-8.0. Then we added Worthington TPCK-treated trypsin (1 mg/mL) dissolved in 1 mM HCl supplemented with 20 mM of CaCl2 to prevent autolysis. The trypsin mixture incubated at 4°C for about 1 hour prior to adding to the protein lysate. Samples were diluted 5-fold by adding 10 mM tris, pH 8.0 to dissolve urea <2M followed by trypsin addition at 1:50 trypsin/protein ratio and samples were incubated overnight at 37°C. After incubation, samples were acidified with TFA to pH 3 or less. Two sequential reverse phase extraction methods were used first. Hydrophilic-Lipophilic Balanced (HLB) was used first and then the flow thru from HLB and wash fractions were vacuum dried, resuspended, and cleaned up again using a C18 solid phase extraction method. The peptides were combined from HLB and C18 cleanups and peptide yield was measured using the BCA peptide assay (Thermo Fisher scientific cat# 23275). Sample digestion efficiency prior to mass spectrometry analysis was inspected by evaluating samples before and after enzymatic digestion using SDS-page gels and again after mass spectrometry analysis. **Phosphoenrichment:** A total of 2 mg of peptides was used for enrichment of phosphopeptides using the Sequential Metal Oxide Affinity Chromatography (SMOAC) Kit (Thermo Fisher Cat# A32993, A32992) protocol. The quantitative analysis of phosphoserine, phosphotyrosine and phosphothreonine peptides by Quantitative Mass Spectrometry was performed as previously described [61, 62] with minor modifications in-tandem using the SMOAC assay. **Mass Spectrometry:** The desalted peptide mixture was fractionated online using EASY-spray columns (25 cm 3 75 mm ID, PepMap RSLC C18 2 mm). The gradient was delivered by an easy-nano Liquid Chromatography 1000 ultra-high-pressure liquid chromatography (UHPLC) system (Thermo Scientific). Tandem mass spectrometry (MS/MS) spectra were collected on the FAIMS TRIBRID mass spectrometer (Thermo Scientific) [63, 64]. Samples were run in biological replicates, and raw MS files were analyzed using MaxQuant (v1.4.1.2) [65]. MS/MS fragmentation spectra were searched using ANDROMEDA against the Uniprot human reference proteome database with canonical and isoform sequences (downloaded August 1^st^ 2021 from http://uniprot.org). N-terminal acetylation, oxidized methionine, and phosphorylated serine, threonine, or tyrosine were set as variable modifications, and carbamidomethyl cysteine (*C) was set as a fixed modification. The false discovery rate was set to 1% using a composite target-reversed decoy database search strategy. Group-specific parameters included max missed cleavages of two and label-free quantitation (LFQ) with an LFQ minimum ratio count of one. Global parameters included match between runs with a match time and alignment time window of 5 and 20 min, respectively, and match unidentified features selected. **Kinase Substrate Enrichment Analysis:** Kinase substrate enrichment analysis (KSEA) Phospho-serine and -threonine peptides were ranked by the signal-to-noise ratio observed for a given perturbation. We then annotated the ranked phospho-peptide lists with predicted upstream kinases from the database of kinase-substrate interactions. For each upstream kinase, we calculated a Kolmogorov–Smirnov score that measured the degree to which its targets were differentially phosphorylated treated samples compared to treatment naïve prostate cancer cell line. This approach is analogous to the normalized enrichment score of gene set enrichment analysis [66] but instead of identifying pathways with a significant fraction of associated members, the procedure identifies kinases with a significant fraction of associated targets. The statistical significance of the KS scores were assessed by permutation analysis. Permutations were performed by randomly shuffling the peptide ranked list, followed by calculation of the KS score for each permutation. After 1,000 permutations, the fraction of randomly ranked lists resulting in a KS score greater than or equal to the observed value was defined as the permutation-based frequency of random occurrence (i.e., the permutation-based p-value). As in GSEA, to normalize for the different number of predictions for each upstream kinase, we calculated the normalized KS score by dividing the observed enrichment score by the mean of the absolute value of all permutation enrichment scores. The false discovery rate for each kinase was calculated using the Benjamini– Hochberg procedure.

### Immunofluorescence

Cells were seeded on coverslips and allowed to attach. Cells were then treated with drugs at the concentrations indicated or DMSO control 1.5 hrs pre irradiation. Cells were irradiated with 1 Gy and harvested at the time points indicated. For γH2AX foci, coverslips were washed 2x5 min in PBS, fixed in 4% paraformaldehyde (ThermoFisher) for 10 min at RT, and then washed 2x5 min in PBS. Coverslips were stored O/N in 3% FBS in PBS at 4°C. Coverslips were then washed 2x5 min in PBS at RT, and permeabilized with 0.5% Triton-X in PBS for 10 min at RT. Coverslips were blocked in 3% serum in PBS (NDS) for 30 min in a moisture chamber and incubated in primary antibody diluted in NDS O/N (γH2AX (1:1000)) at 4°C. Coverslips were washed 3x10 min with PBS, incubated with secondary antibody conjugated with Texas-Red in 1% NDS for 1 hr at RT and washed 3x10 min at RT. The coverslips were then mounted and sealed with clear nail polish. All coverslips were stored at 4°C until imaged. Images were acquired with a Lionheart™ FX Automated Microscope and raw images were quantified with Imaris spot counter module (v10.0.1). Selected images for publication were pre-processed for background flattening with consistent rolling ball diameter and subsequently deconvoluted using Gen5 software (v3.11).

### DNA Fiber Assays

For experiments using VX-970 and M3814, C4-2 ATM KO19 cells were plated 48-72 hours before to ensure that plated cells were actively dividing. Cells were treated with DMSO control, 1 μM VX-970, 10 μM M3814, or combination 1.5 hrs before pulse-labeling to allow kinase inhibitor integration into the cells. For Compound B/C experiments, C4-2 ATM KO19 cells were treated with 100 nM of either compound for 72 hours before pulse-labeling. Exponentially growing cells were pulse-labeled for 20 minutes with 25 µM 5-iodo-2-deoxyuridine (I7125, Sigma-Aldrich), followed by a second 20-minute pulse with 250 µM 5-chloro-2-deoxyuridine (C6891, Sigma-Aldrich). The labeled cells were then washed twice with ice-cold 1X PBS, collected, and suspended at a concentration of 30,000 cells/ml. Subsequently, 30 µl of the suspension was centrifuged onto slides for 4 minutes at 800 rpm. After cytospinning, the slides were immersed in Lysis Buffer (0.5% SDS, 200mM Tris-HCl, 50mM EDTA) for 5 minutes, and DNA molecules were stretched using a homemade LEGO device. DNA fiber spreads were fixed in ice-cold Carnoy fixative for 10 minutes at room temperature and air-dried. Slides were rehydrated twice in water and incubated for 1 hour at room temperature in 2.5 M HCl. Afterward, the slides were rinsed twice in 1X PBS and blocked for 1 hour at room temperature in a blocking solution (1X PBS + 1% BSA + 0.5% Triton X-100 + 0.02% NaN3). The slides were then incubated in primary antibodies overnight at 4°C. The following primary antibodies were used at the indicated dilutions: 1:100 anti-BrdU (BDB347580, Becton Dickson) and 1:250 anti-CldU (ab6326, Abcam). The slides were then rinsed three times in 1X PBS and fixed in 4% paraformaldehyde in PBS for 10 minutes at room temperature. Afterward, they were rinsed twice in 1X PBS and incubated with 1:1,000 dilutions of Alexa Fluor-conjugated donkey anti-mouse or donkey anti-rat secondary antibodies (Invitrogen) for 2 hours at room temperature. Finally, the slides were washed twice in 1X PBS and mounted in ProLong Gold antifade mounting solution. Immunofluorescence images were captured on a DeltaVision Ultra (Cytiva) microscope system equipped with a 4.2 Mpx sCMOS detector. Fibers were acquired with a x60 objective (PlanApo N 1.42 oil) and 1 × 0.2 μm z-section. Quantitative image analyses were performed using Fiji (v.2.1.0/1.53c). IdU and CldU track lengths were measured using the line tool. Only intact fibers incorporating both IdU and CldU were measured.

### Animal Experiments

All animals were housed and studies conducted at UTSW in a sterile facility. All animal work was done under the supervision of the Animal Resource Center (ARC) and approved by the Institutional Animal Care and Use Committee (IACUC). For studies with ATM-proficient lines, 4X10^6^ 22Rv1 V2 cells were injected subcutaneously into the R flank of male nude mice in 50% ice cold PBS and 50% matrigel (Corning). For studies with ATM-deficient cell lines, 6X10^6^ 22Rv1 KO5 cells were injected subcutaneously into the R flank of male Crl:NU(NCr)-Foxn1^nu^ (nude) mice in 50% ice cold PBS and 50% matrigel (Corning). When tumor volume reached ∼200 mm^3^, mice were randomized. The treatment schedule was as follows: for mice without irradiation, mice were treated twice daily with 100 ccs 62.5 mg/kg (125 mg/kg/day) Compound B dissolved in PEG200, or twice daily PEG200 via oral gavage with flexible gavage tips (Instech) for 6 days. For mice receiving irradiation, mice were treated twice daily with 62.5 mg/kg (125 mg/kg/day) Compound B dissolved in PEG200 or twice daily PEG200 control for 6 days. Targeted radiation (3 Gy) directed to the tumor was given 4 hours after the first dose on days 2 and 4. Mice were given moist chow for 14 days starting on treatment day 1, regardless of treatment condition and were calipered and weighed periodically.

Compound B was dissolved via prolonged vortexing of the drug (∼30 min) and heating to 37°C for 15 min.

### Data Accessibility

The raw phosphoproteomic data in this manuscript can be accessed with accession number # PXD050955 at https://massive.ucsd.edu/ProteoSAFe/index.jsp.

## Key Resources and Materials Table

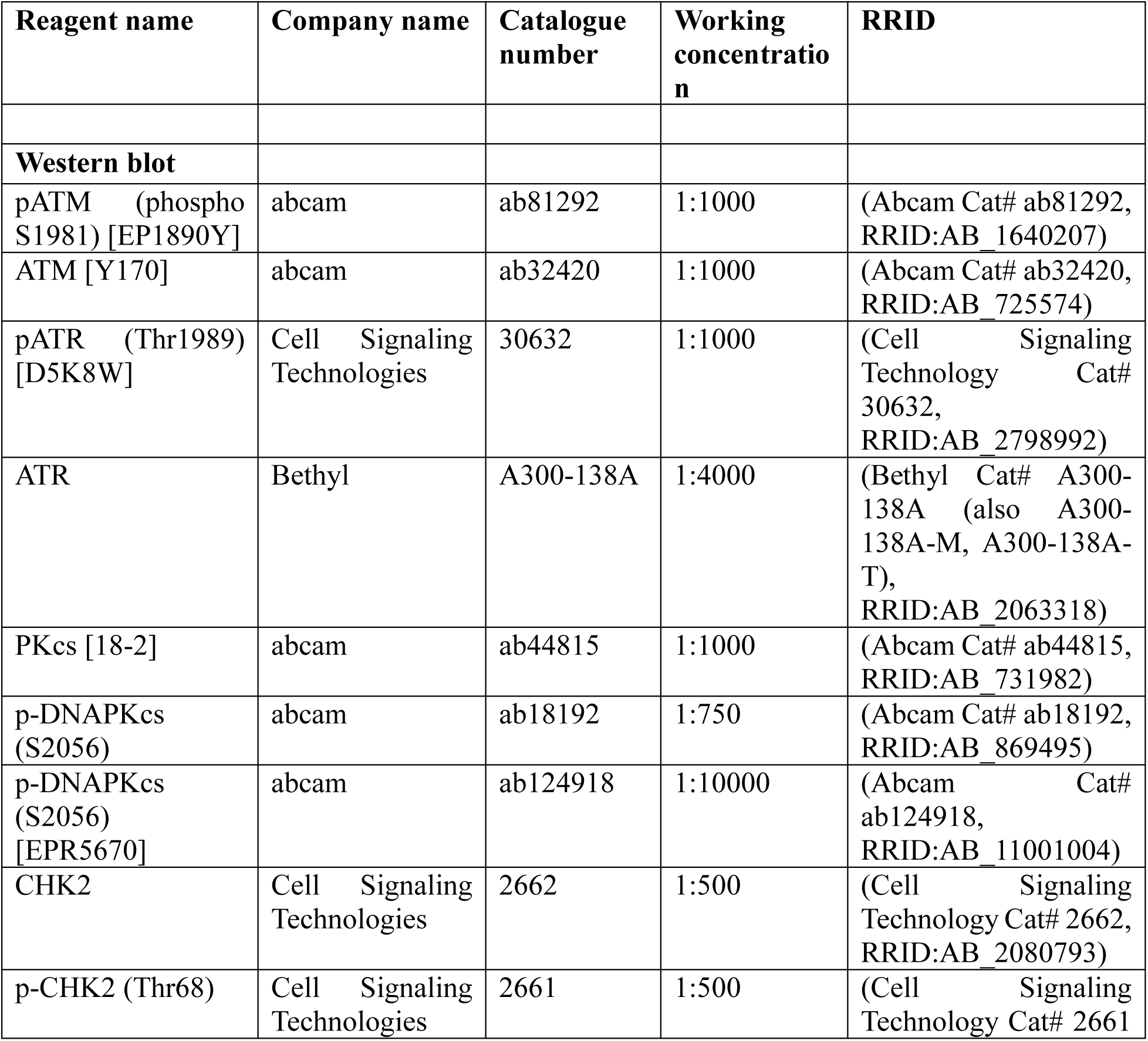
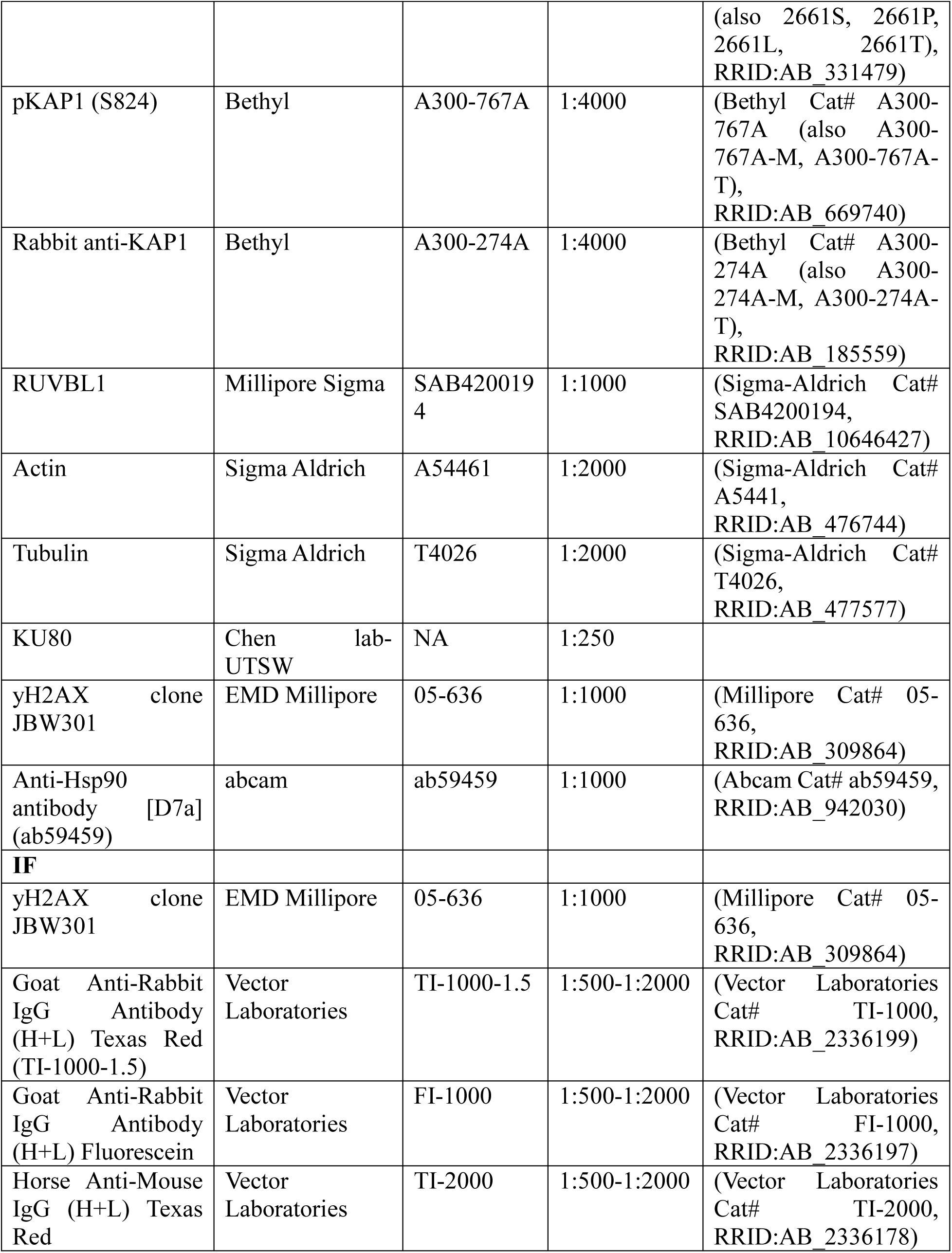
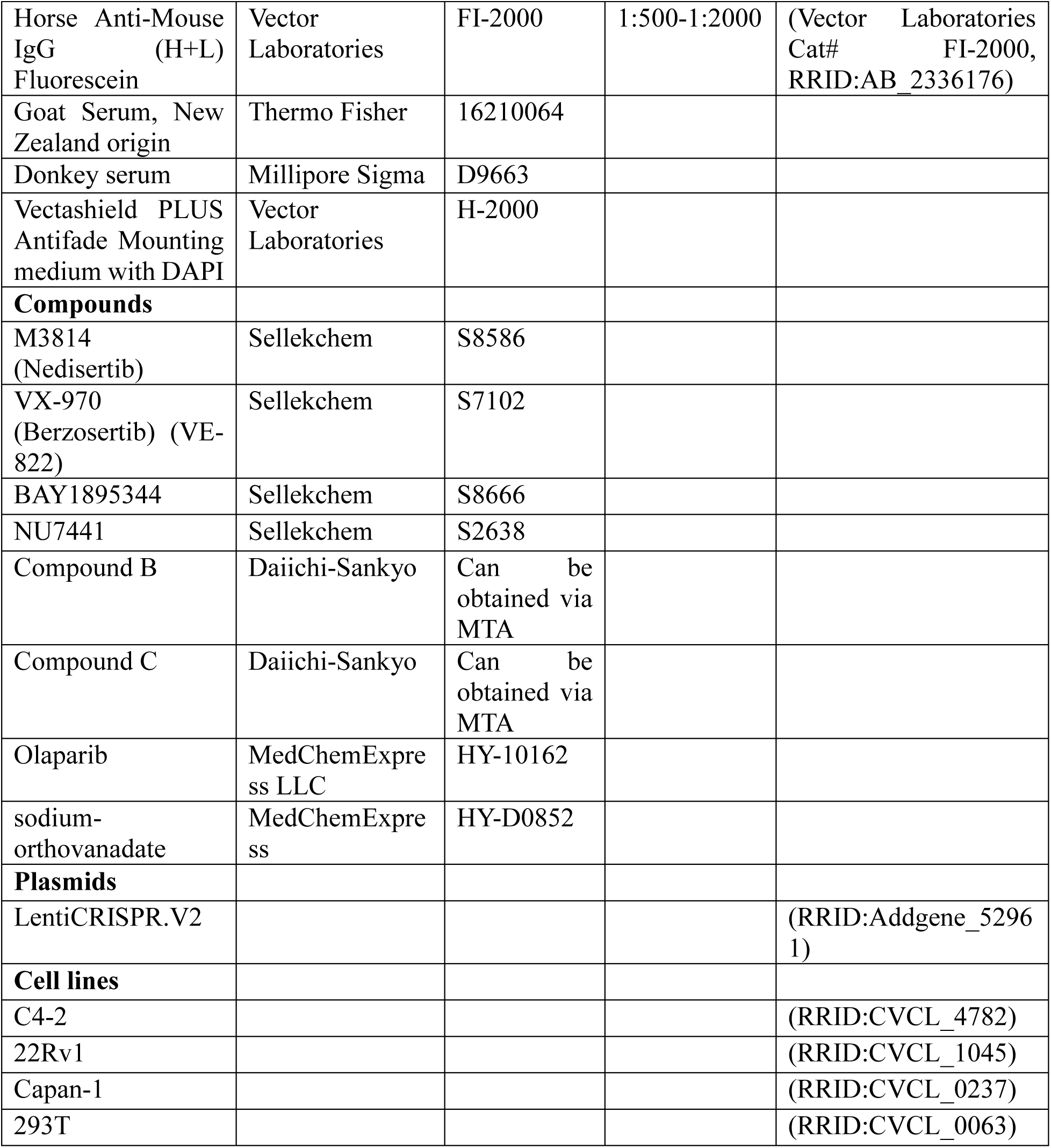

## Acknowledgements

We would like to thank Dr. Ganesh Raj for his guidance and oversight on this project. We would also like to thank Dr. Kathryn O’Donnell for her support and mentorship throughout this project.

---

**Supplemental Figure S1.**
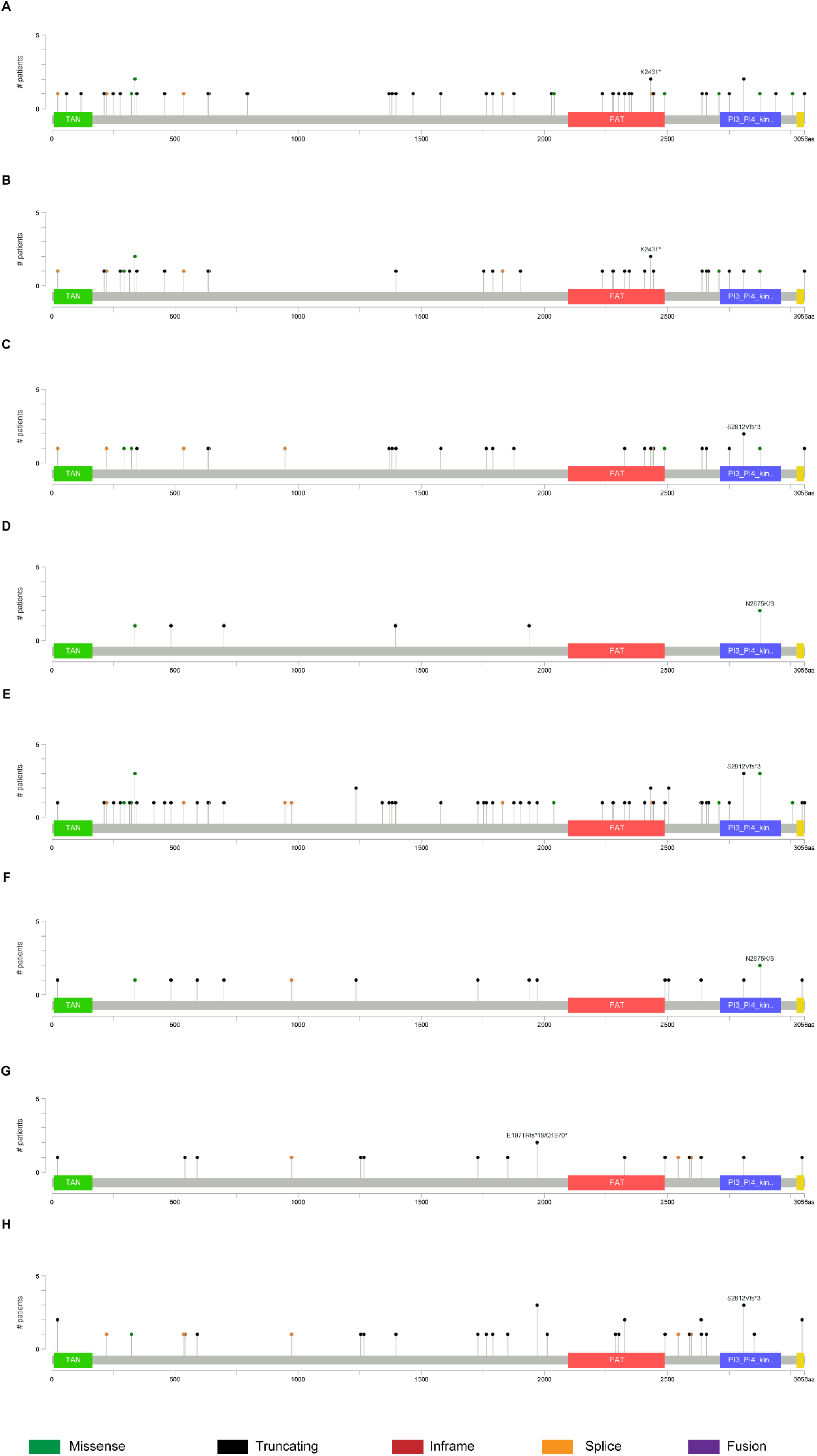
Lollipop graphs of multiple independent datasets demonstrate that pathogenic ATM mutations are often truncating and are distributed throughout the genome. All lollipop graphs were generated with cBioPortal on 1.15.24. All include both germline and somatic mutations and excluded variants of unknown significance. **S1a** Lollipop graph generated from Race Differences in Prostate Cancer dataset [21]. N=2069 samples/patients found 53 pathogenic ATM mutations. 36/53 (67.9%) are truncation mutations. **S1b** Lollipop graph generated from MSK Clinical Cancer Research dataset [29]. N=1417 samples/patients found 38 pathogenic ATM mutations. 25/38 (65.7%) are truncation mutations. **S1c** Lollipop graph generated from MSK European Urology dataset [30]. N=1465 samples/patients found 32 pathogenic ATM mutations. 20/32 (62.5%) are truncation mutations. **S1d** Lollipop graph generated from the TCGA dataset [28]. N=489 samples/patients found 9 pathogenic ATM mutations. 4/9 (44.4%) are truncation mutations. **S1e** Lollipop graph generated from all prostate adenocarcinomas on cBioPortal [20, 23–27, 29–32, 34]. N= 5675 samples from 5554 patients found 106 pathogenic ATM mutations. 70/106 (66.0%) are truncation mutations. **S1f** Lollipop graph of MSK/DFCI prostate adenocarcinoma dataset [20]. N= 1013 samples/patients found 17 pathogenic ATM mutations. 13/17 (76.5%) are truncation mutations. **S1g** Lollipop graph of metastatic patients in the SU2C/PCF Dream Team dataset [67]. N= 444 samples from 429 patients found 21 pathogenic ATM mutations. 18/21 (85.7%) are truncation mutations. **S1h** Lollipop graph of all metastatic prostate cancer datasets [10, 19, 22, 67]. N=1139 samples from 1122 patients found 41 pathogenic ATM mutations. 34/41 (82.9%) are truncation mutations.

**Supplemental Figure S2.**
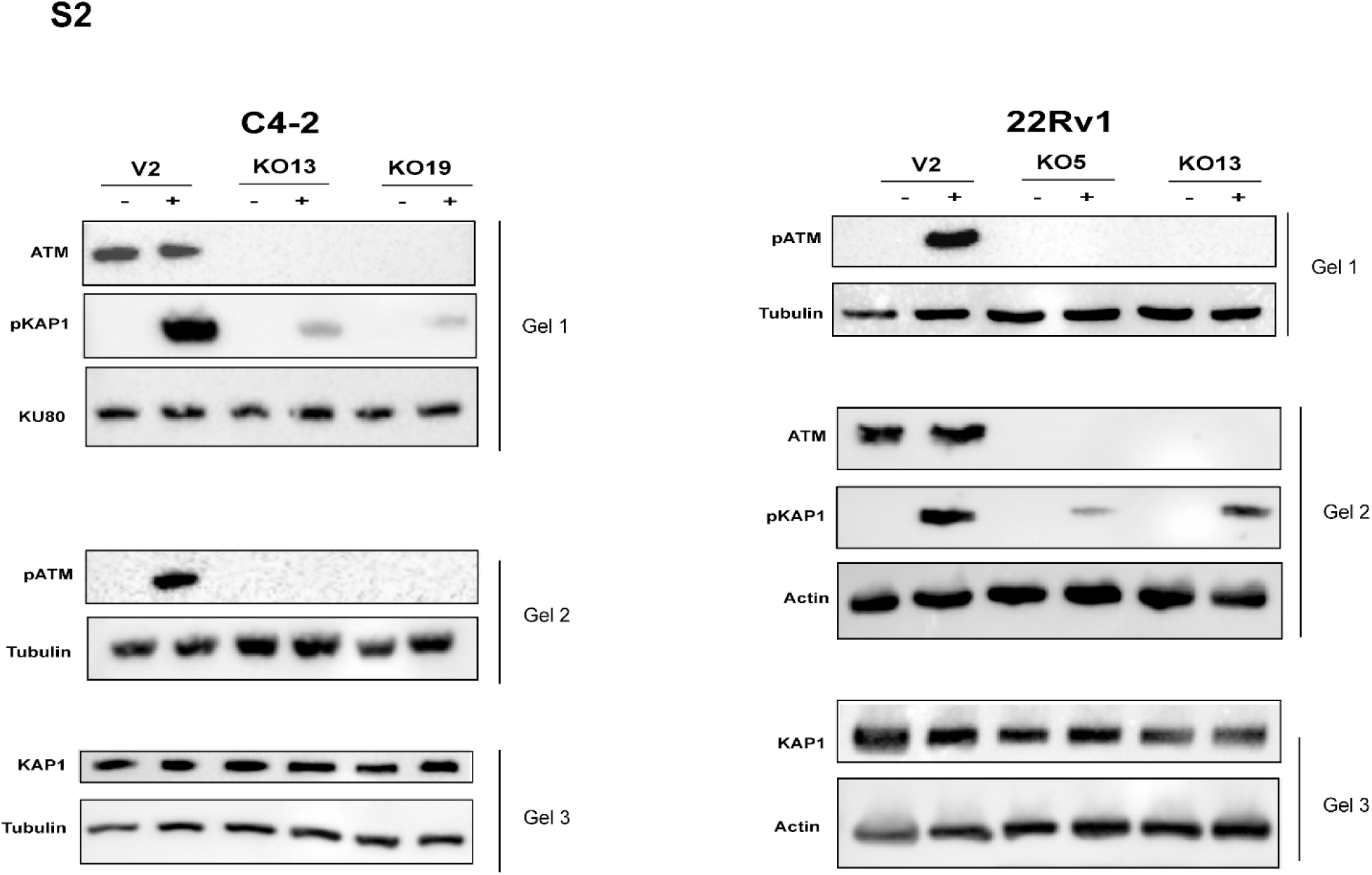
C4-2 and 22Rv1 V2 and KO lines were run on three separate blots each. The same lysates were used for each blot. Each respective blot with its loading control is shown. For ease of presentation, ***Figure 1c*** and ***Figure 1d*** only show one loading control.

**Supplemental Figure S3.**
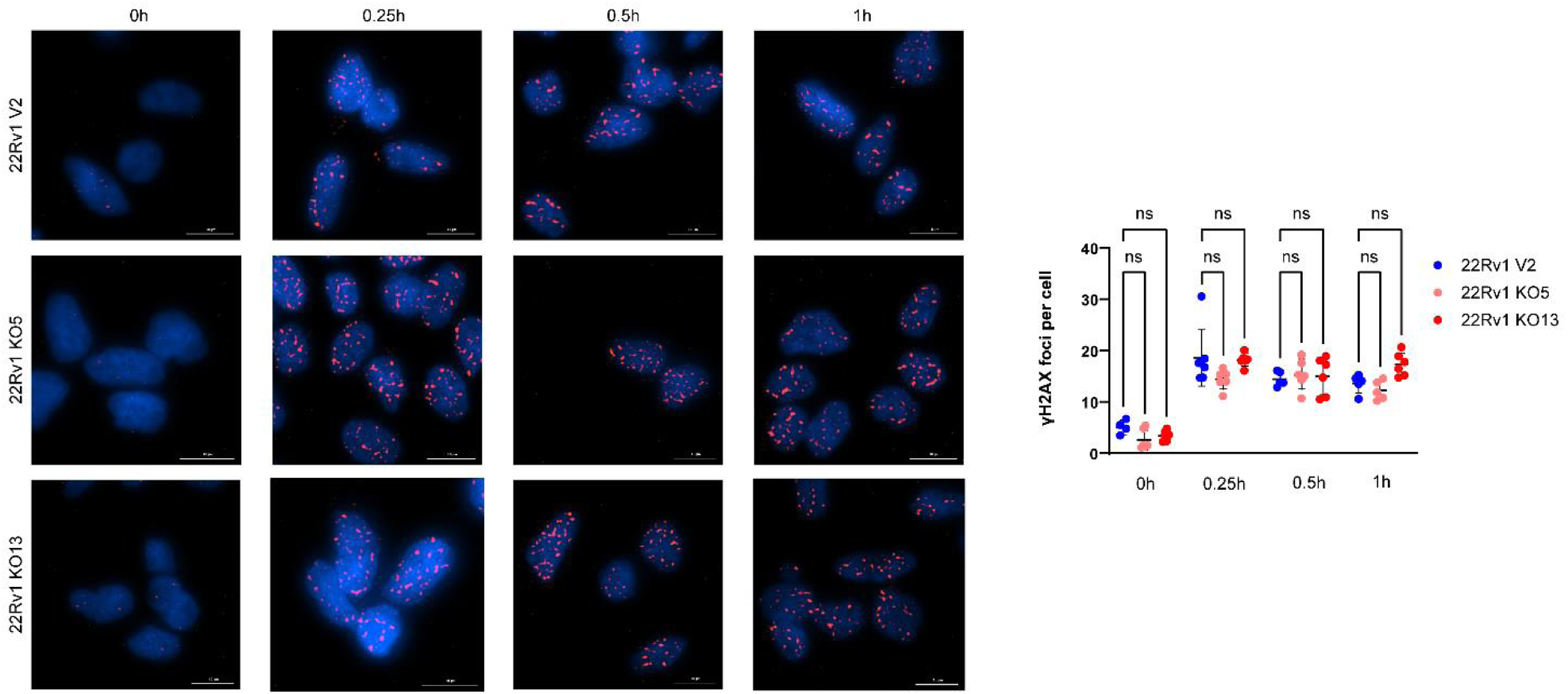
**S3** Time course IF staining of γH2AX foci recruitment in response to 1 Gy IR on DMSO treated 22Rv1 V2, KO5, and KO13 cells demonstrates no statistically significant difference in number of γH2AX foci recruited up to 1 hr after IR. Two independent experiments were run; data presented is from one experiment. A minimum of 60 cells were quantified per condition in the selected experiment. Representative images were pre-processed for background flattening with consistent rolling ball diameter and subsequently deconvoluted using Gen5 software. Scale bar represents 10 µM. Foci were quantified using the Spot Counting module on Imaris software. Each dot represents an image field. Statistical significance was calculated with One-Way ANOVA with Tukey’s multiple comparison test.

**Supplemental Figure S4.**
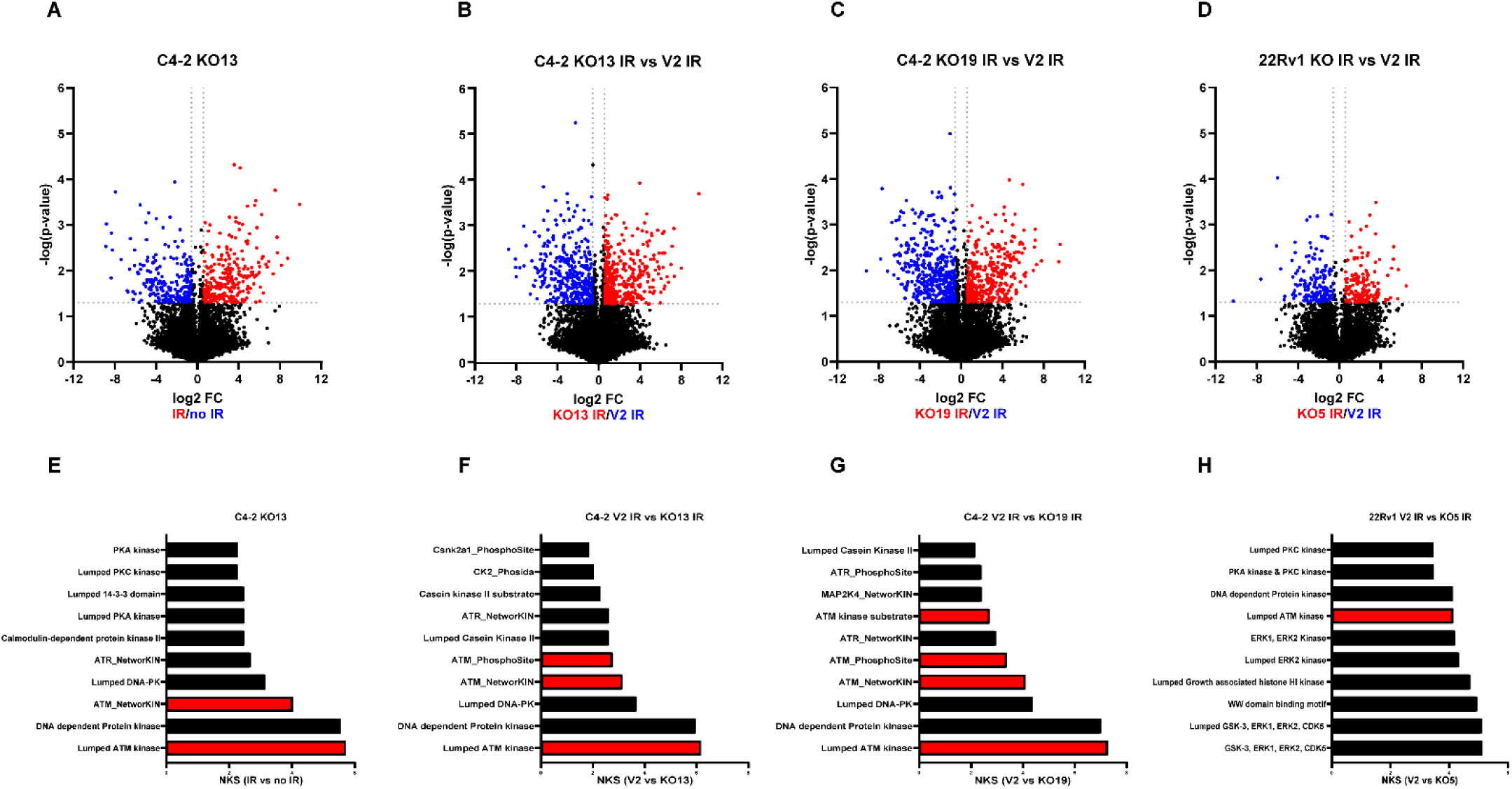
**S4a-d** Volcano plot of C4-2 KO13 IR vs no IR **(S4a),** Volcano plot of C4-2 KO13 IR vs C4-2 V2 IR **(S4b)** Volcano plot of C4-2 KO19 IR vs C4-2 V2 IR **(S4c)** and Volcano plot of 22Rv1 KO5 IR vs 22Rv1 V2 IR (**S4c)** demonstrate distinct panels of differentially expressed phosphopeptides in response to IR. Volcano plots were generated with Quantum-Normalized Log Transformed (QNLT) values. Vertical dotted line represents fold change cut off of 1.5. Horizontal dotted line represents p-value 0.05. **S4e** Kinase Substrate Enrichment Analyses (KSEA) of C4-2 KO13 presented as irradiated samples/non-irradiated controls. The pathways were filtered for number of hits > 5 and FDR < 0.05. The top ten pathways, by Normalized Kolmogorov–Smirnov score (NKS score), are shown. Analyses demonstrate that despite convincing loss of ATM in C4-2 ATM KO13, there is still a dependence on the ATM pathway. **S4f-h** KSEA of C4-2 V2 IR vs C4-2 KO13 IR **(S4f)**, C4-2 V2 IR vs C4-2 KO19 IR **(S4g)**, and 22Rv1 V2 IR vs 22Rv1 KO5 IR (**S4h)** KSEA demonstrates that ATM pathways are upregulated in V2 cells over ATM KO lines, further confirming ATM kinase attenuation in these cell lines.

**Supplemental Figure S5.**
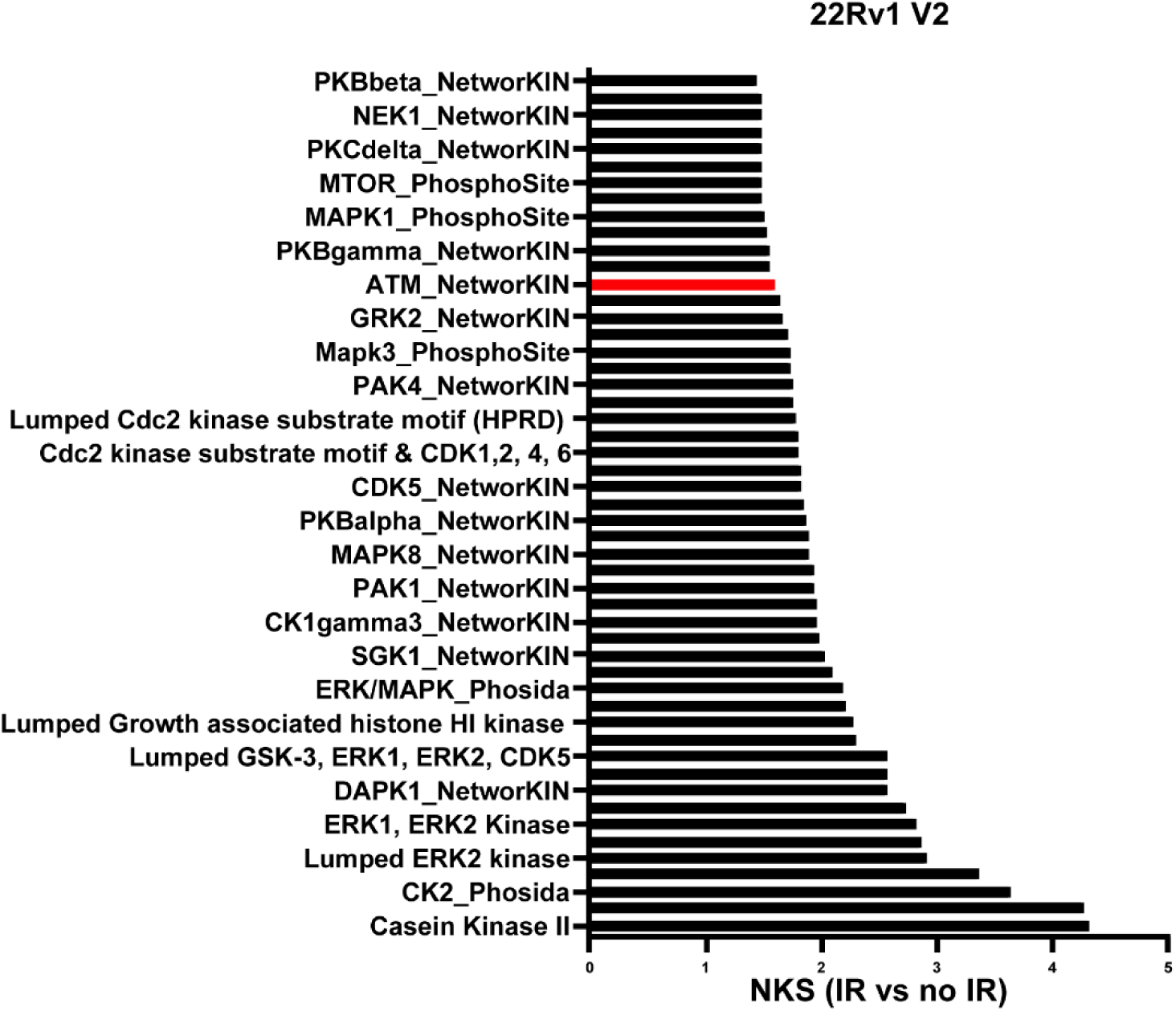
KSEA of 22Rv1 V2 presented as irradiated samples/non-irradiated controls. The pathways were filtered for number of hits > 5 and FDR < 0.05. The top pathways, by Normalized Kolmogorov–Smirnov score (NKS score), are shown. ATM_KIN is a significantly enriched pathway in response to IR in this cell line.

**Supplemental Figure S6.**
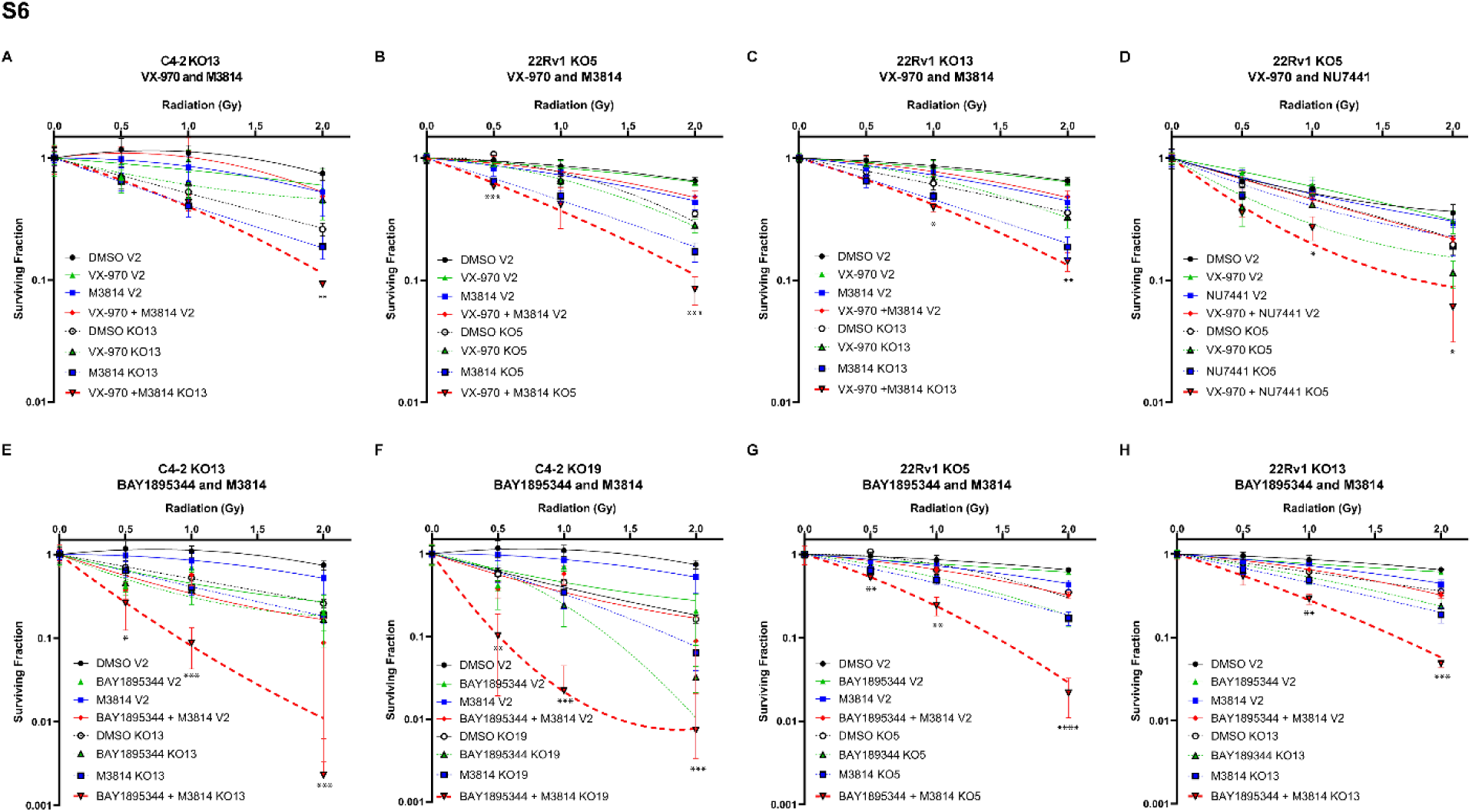
**S6a** Clonogenic survival assay of C4-2 V2 and C4-2 KO13 with 200 nM M3814 and 10 nM VX-970 and escalating doses of irradiation demonstrates that combination therapy is most effective in the context of ATM-deficiency. Note that this experiment was in conjunction with **Figure 3a**, so the C4-2 V2 control curve is consistent. **S6b and S6c** Clonogenic survival assay of 22Rv1 V2 and 22Rv1 KO5 **(S6b)** and 22Rv1 KO13 **(S6c)** with 100 nM M3814 and 10 nM VX-970 and escalating doses of irradiation demonstrates that combination therapy is most effective in the context of ATM loss **S6d** Clonogenic survival assay of 22Rv1 V2 and 22Rv1 KO5 with 200 nM of a different DNA-PKcs inhibitor, NU7441, and 10 nM VX-970 demonstrates combination therapy is most effective in preventing DDR; independent of the DNA-PKcs inhibitor used. **S6e and S6f** Clonogenic survival assay of C4-2 KO13 **(S6e)** and C4-2 KO19 **(S6f)** compared to C4-2 V2 with 200 nM M3814, and a 10 nM of a different ATR inhibitor, BAY1895344, demonstrates a similar pattern, with combination therapy being most effective in preventing DDR; independent of the ATR inhibitor used. **S6g and S6h** Clonogenic survival assay of 22Rv1 KO5 **(S6g)** and 22Rv1 KO13 **(S6h)** compared to 22Rv1 V2 with 100 nM M3814 and 5 nM of a different ATR inhibitor BAY1895344 demonstrates a similar pattern, with combination therapy being most effective in preventing DDR; independent of the ATR inhibitor used. For **S6a-h** all values represent mean ± SD in technical triplicate. All colonies were quantified with ImageJ cell counter plugin, and survival curve was generated using the linear quadratic equation (Y is fraction of cells surviving) on GraphPad Prism. Please note that **S6a, S6e, and S6f** were plated concurrently and therefore share a C4-2 V2 control curve, and **S6b, S6c, S6g, and S6h** were plated concurrently and share a 22Rv1 V2 control curve. All statistics shown analyze KO DMSO condition vs. KO combination therapy condition; determined by multiple t-tests with the Holm-Šídák method

**Supplemental Figure S7.**
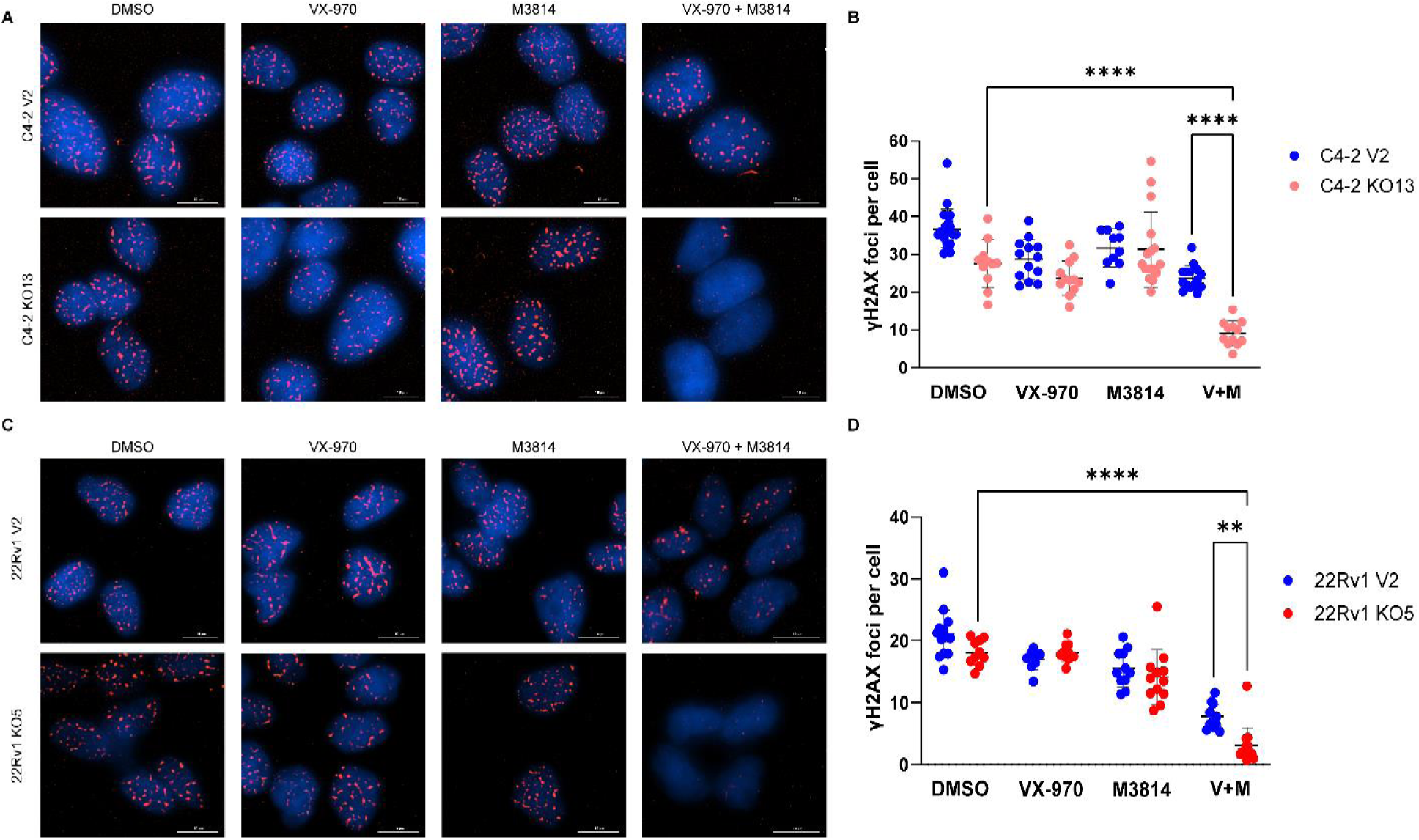
**S7a-b** IF staining of C4-2 V2 and C4-2 KO13 cells for γH2AX foci. Cells were pre-treated with DMSO control, 1 µM VX-970, 10 µM M3814, or 1 µM VX-970 + 10 µM M3814 for 1.5 hrs, subjected to 1 Gy IR, and harvested 1 hr later. γH2AX foci are significantly decreased at 1 hr in the dual therapy group. Scale bar represents 10 µm. Data presented is from one experiment, with a minimum of 58 cells counted per condition per cell line. Representative images were pre-processed for background flattening with consistent rolling ball diameter and subsequently deconvoluted using Gen5 software. Note that this experiment was run concurrently with ***Figure 3b-c***, so the control C4-2 V2 pictures and values remain consistent. γH2AX foci were quantified using the Spot Counting module on Imaris software. Each dot represents an image field. Error bars represent mean ±SD. Quantification of γH2AX foci per cell demonstrates a statistically significant reduction in combination therapy in KO19 as opposed to DMSO control. An Ordinary One Way ANOVA with Turkey’s multiple comparison test was used for statistical analysis. (**** represents p-value <0.0001). **S7c-d** IF staining of 22Rv1 V2 and 22Rv1 KO5 cells for γH2AX foci. Cells were treated as in C4-2 condition. γH2AX foci are significantly decreased at 1 hr in the dual therapy group. Scale bar represents 10 µm. Two independent experiments were run; data presented is from one experiment. Representative images were pre-processed for background flattening with consistent rolling ball diameter and subsequently deconvoluted using Gen5 software (**S7c)**. γH2AX foci were quantified using the Spot Counting module on Imaris software. A minimum of 70 cells were quantified per condition in the selected experiment. Each dot represents an image field. Error bars represent mean ± SD (**S7d**). Quantification of γH2AX foci per cell demonstrates a statistically significant reduction in combination therapy as opposed to DMSO control in 22Rv1 KO5. An Ordinary One Way ANOVA with Turkey’s multiple comparison test was used for statistical analysis.

**Supplemental Figure S8.**
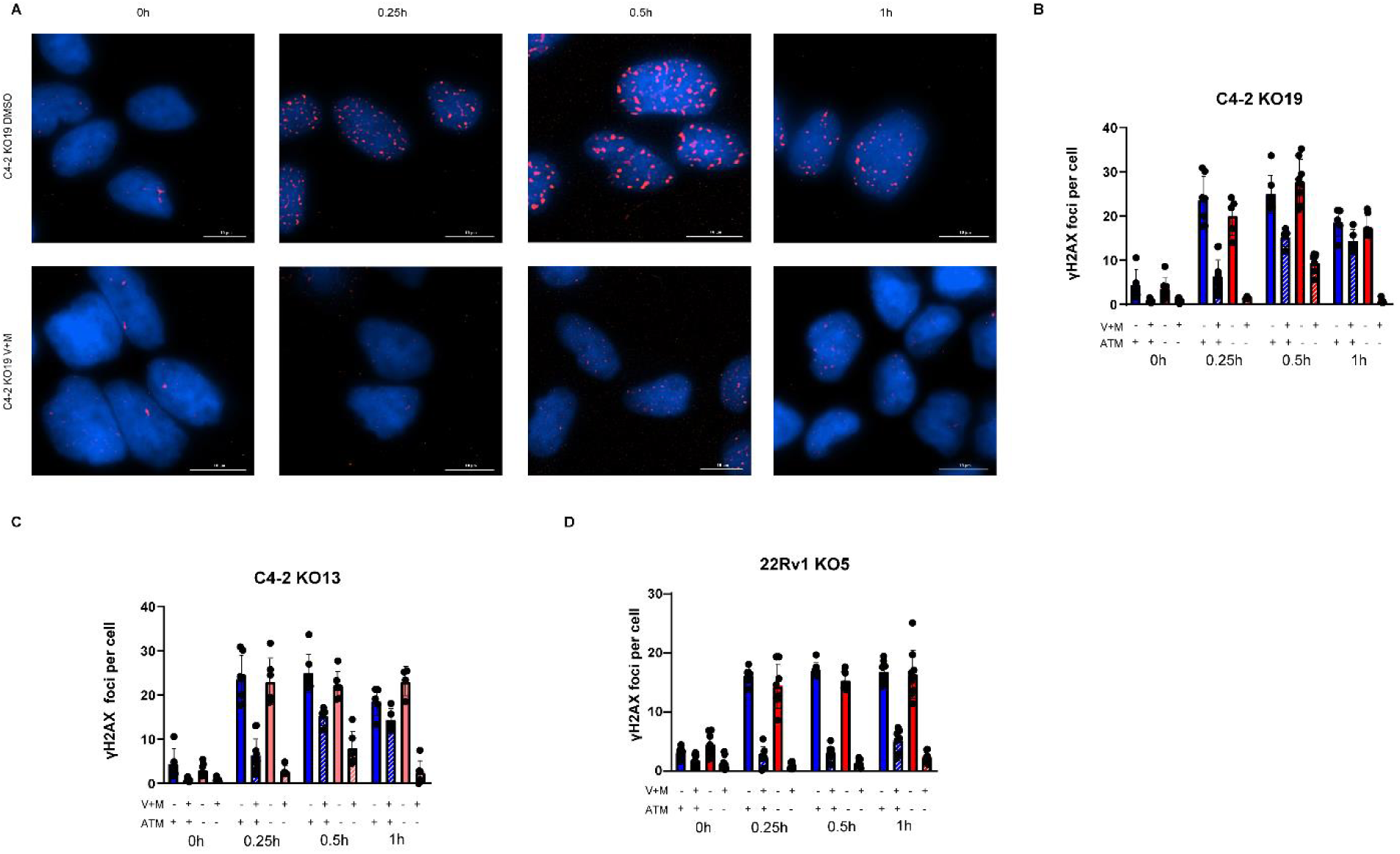
**S8a-c** IF staining of C4-2 V2 and C4-2 KO19 **(S8a-b)** and C4-2 KO13 **(S8c)** cells for γH2AX foci demonstrates that the decrease of γH2AX foci at 1 hr with combination therapy in ATM-deficient cells is due to an inability to recruit foci. Cells were pre-treated with DMSO control or 1 µM VX-970+ 10 µM M3814 for 1.5 hrs, subjected to 1 Gy IR, and harvested at the time points indicated. Scale bar represents 10 µm. Data presented is from one experiment, with a minimum of 59 cells counted per condition per cell line. Representative C4-2 KO19 images **(S8a)** were pre-processed for background flattening with consistent rolling ball diameter and subsequently deconvoluted using Gen5 software. Error bars represent mean ±SD. **S8d** IF staining of 22Rv1 V2 and 22Rv1 KO5 for γH2AX foci demonstrates that the lack of γH2AX foci at 1 hr with combination therapy in ATM-deficient cells is due to an inability to recruit foci. Conditions are the same as described for C4-2 cells. Data presented is from one independent experiment with a minimum of 66 cells quantified per condition. Error bars represent mean ±SD.

**Supplemental Figure S9.**
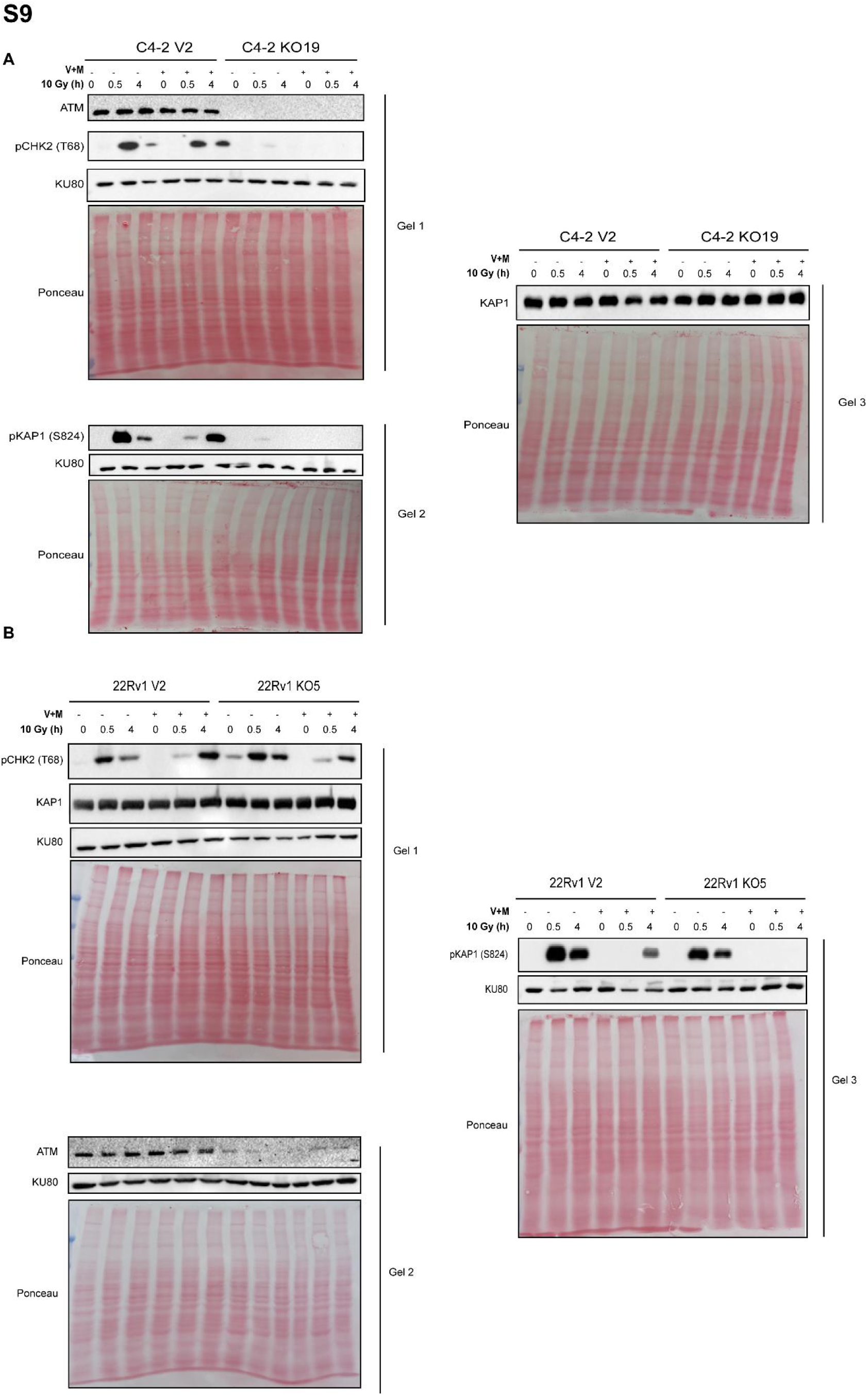
C4-2 and 22Rv1 V2 and KO lines were run on multiple blots. The same lysates were used for each blot. Each respective blot with its loading control is shown here. For ease of presentation, **Figure 3e** and **Figure 3f** only show one loading control.

**Supplemental Figure S10.**
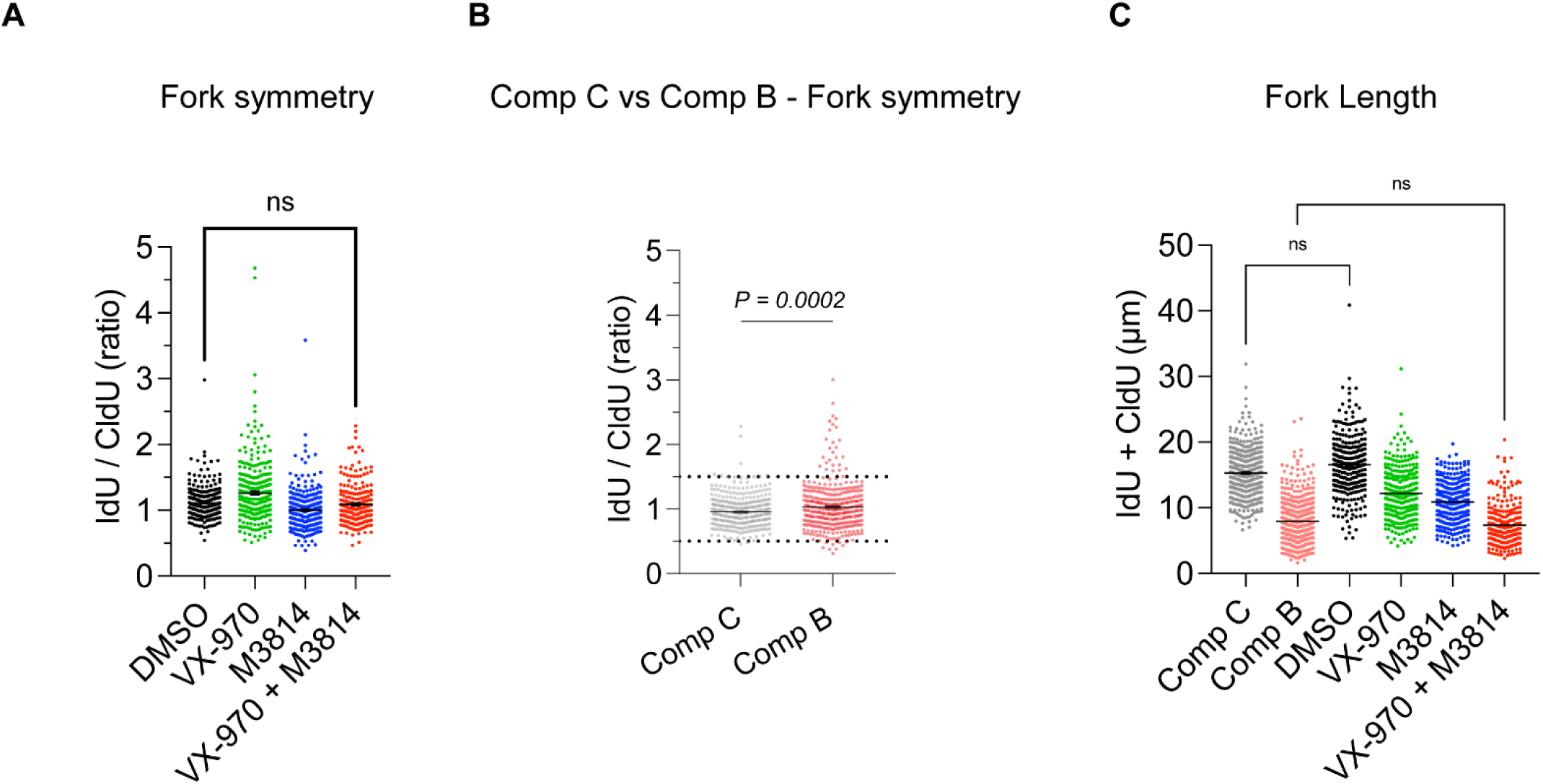
**S10a** C4-2 ATM KO19 cells were treated with either DMSO control, 1 μM VX-970, 10 μM M3814, or VX-970+M3814 for 1.5 hours and subsequently harvested. Fiber assay analysis demonstrate no difference in fork symmetry between DMSO control and dual treated C4-2 KO19 cells. Data are mean ± S.E.M. from DMSO, n=300; VX-970, n= 300, M3814, n= 301; VX-970 + M3814, n=234 fibers pooled from 2 independent experiments. Statistics calculated by Kruskal-Wallis multiple comparison test. **S10b** C4-2 KO19 cells were treated with either 100 nM Compound B or 100 nM Compound C for 72 hours and subsequently harvested. Minimal difference is seen in fork symmetry between Compound B and Compound C control. Data are mean ± S.E.M from n=450 (Compound B) and n=451 (Compound C) forks pooled from 3 independent experiments. Statistics were calculated with Mann-Whitney U test (unpaired and nonparametric). **S10c** Combination of data for both dual therapy and Compound B/C treatment demonstrates no statistical difference in fork length between DMSO and Compound C or Compound B and VX-970+M3814 treatment. Values are presented as mean ± S.E.M; statistics calculated by Kruskal-Wallis test with Dunn’s multiple comparisons.

**Supplemental Figure S11.**
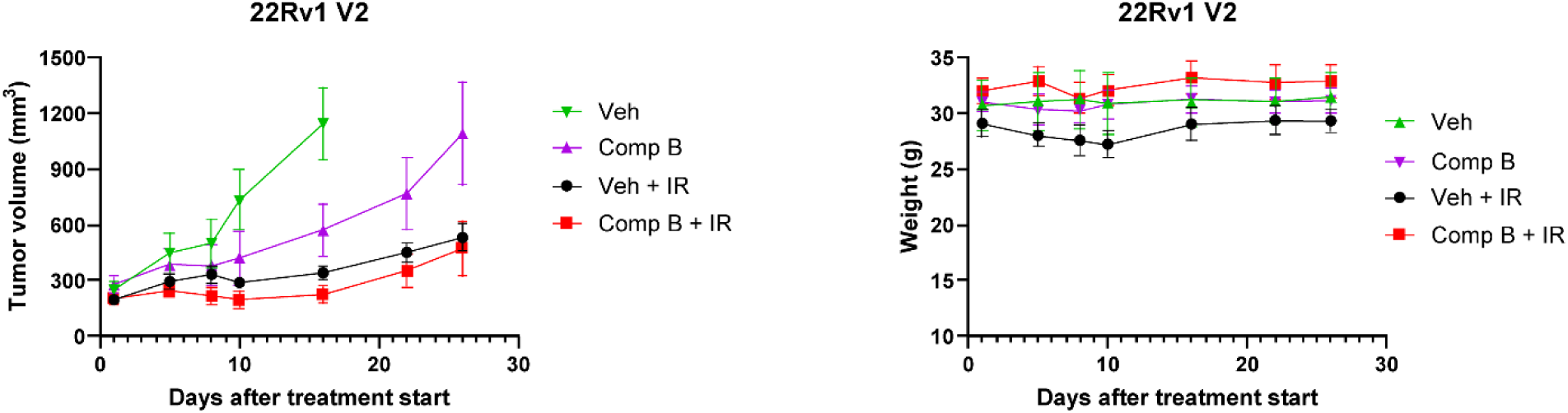
*In vivo* animal experiments with ATM-proficient cells. 22Rv1 V2 cells were xenografted into male nude mice and when tumor volume reached ∼200 mm^3^ mice were randomized. Mice were treated ± Compound B for 6 days, with or without 3 Gy IR on days 2 and 4. Schematic of treatment outlined in ***Figure 5j***. Raw tumor volumes and mouse weight were monitored, and are presented as mean ± S.E.M. No statistical significance was noted between vehicle + IR and Compound B+ IR using a 2-Way ANOVA with Bonferroni’s multiple comparison’s test. Vehicle n=4, Compound B n=5, Vehicle + IR n=5, Compound B + IR n=4.

## References

1. Siegel, R.L., et al., Cancer statistics, 2023. CA: A Cancer Journal for Clinicians, 2023. 73(1): p. 17–48.

2. Eastham, J.A., et al., Clinically Localized Prostate Cancer: AUA/ASTRO Guideline, Part I: Introduction, Risk Assessment, Staging, and Risk-Based Management. J Urol, 2022. 208(1): p. 10–18.

3. Lowrance, W., et al., Updates to Advanced Prostate Cancer: AUA/SUO Guideline (2023). J Urol, 2023. 209(6): p. 1082–1090.

4. de Bono, J.S., et al., Abiraterone and Increased Survival in Metastatic Prostate Cancer. New England Journal of Medicine, 2011. 364(21): p. 1995–2005.

5. Scher, H.I., et al., Antitumour activity of MDV3100 in castration-resistant prostate cancer: a phase 1-2 study. The Lancet, 2010. 375(9724): p. 1437–1446.

6. Tran, C., et al., Development of a Second-Generation Antiandrogen for Treatment of Advanced Prostate Cancer. Science, 2009. 324(5928): p. 787–790.

7. Hofstad, M., et al., Alterations in BRCA2 as Determinants of Therapy Response in Prostate Cancer. Critical Reviews™ in Oncogenesis, 2022. 27(1).

8. Buttigliero, C., et al., Understanding and overcoming the mechanisms of primary and acquired resistance to abiraterone and enzalutamide in castration resistant prostate cancer. Cancer Treatment Reviews, 2015. 41(10): p. 884–892.

9. Mekonnen, N., H. Yang, and Y.K. Shin, Homologous Recombination Deficiency in Ovarian, Breast, Colorectal, Pancreatic, Non-Small Cell Lung and Prostate Cancers, and the Mechanisms of Resistance to PARP Inhibitors. Frontiers in Oncology, 2022. 12.

10. Robinson, D., et al., Integrative clinical genomics of advanced prostate cancer. Cell, 2015. 161(5): p. 1215–1228.

11. de Bono, J., et al., Olaparib for Metastatic Castration-Resistant Prostate Cancer. N Engl J Med, 2020. 382(22): p. 2091–2102.

12. Marshall, C.H., et al., Differential Response to Olaparib Treatment Among Men with Metastatic Castration-resistant Prostate Cancer Harboring BRCA1 or BRCA2 Versus ATM Mutations. Eur Urol, 2019. 76(4): p. 452–458.

13. Stopsack, K.H., Efficacy of PARP Inhibition in Metastatic Castration-resistant Prostate Cancer is Very Different with Non-BRCA DNA Repair Alterations: Reconstructing Prespecified Endpoints for Cohort B from the Phase 3 PROfound Trial of Olaparib. Eur Urol, 2020.

14. Anscher, M.S., et al., FDA Approval Summary: Rucaparib for the Treatment of Patients with Deleterious BRCA-Mutated Metastatic Castrate-Resistant Prostate Cancer. Oncologist, 2021. 26(2): p. 139–146.

15. Chi, K.N., et al., Niraparib and Abiraterone Acetate for Metastatic Castration-Resistant Prostate Cancer. J Clin Oncol, 2023. 41(18): p. 3339–3351.

16. Abida, W., et al., Rucaparib in Men With Metastatic Castration-Resistant Prostate Cancer Harboring a BRCA1 or BRCA2 Gene Alteration. J Clin Oncol, 2020. 38(32): p. 3763–3772.

17. Fizazi, K., et al., Rucaparib or Physician’s Choice in Metastatic Prostate Cancer. New England Journal of Medicine, 2023. 388(8): p. 719–732.

18. Yenerall, P., et al., RUVBL1/RUVBL2 ATPase Activity Drives PAQosome Maturation, DNA Replication and Radioresistance in Lung Cancer. Cell Chem Biol, 2020. 27(1): p. 105–121.e14.

19. Grasso, C.S., et al., The mutational landscape of lethal castration-resistant prostate cancer. Nature, 2012. 487(7406): p. 239–43.

20. Armenia, J., et al., The long tail of oncogenic drivers in prostate cancer. Nat Genet, 2018. 50(5): p. 645–651.

21. Stopsack, K.H., et al., Differences in Prostate Cancer Genomes by Self-reported Race: Contributions of Genetic Ancestry, Modifiable Cancer Risk Factors, and Clinical Factors. Clin Cancer Res, 2022. 28(2): p. 318–326.

22. Stopsack, K.H., et al., Oncogenic Genomic Alterations, Clinical Phenotypes, and Outcomes in Metastatic Castration-Sensitive Prostate Cancer. Clin Cancer Res, 2020. 26(13): p. 3230–3238.

23. Baca, S.C., et al., Punctuated evolution of prostate cancer genomes. Cell, 2013. 153(3): p. 666–77.

24. Barbieri, C.E., et al., Exome sequencing identifies recurrent SPOP, FOXA1 and MED12 mutations in prostate cancer. Nat Genet, 2012. 44(6): p. 685–9.

25. Fraser, M., et al., Genomic hallmarks of localized, non-indolent prostate cancer. Nature, 2017. 541(7637): p. 359–364.

26. Kumar, A., et al., Substantial interindividual and limited intraindividual genomic diversity among tumors from men with metastatic prostate cancer. Nat Med, 2016. 22(4): p. 369–78.

27. Taylor, B.S., et al., Integrative genomic profiling of human prostate cancer. Cancer Cell, 2010. 18(1): p. 11–22.

28. Hoadley, K.A., et al., Cell-of-Origin Patterns Dominate the Molecular Classification of 10,000 Tumors from 33 Types of Cancer. Cell, 2018. 173(2): p. 291–304.e6.

29. Chakraborty, G., et al., The Impact of PIK3R1 Mutations and Insulin-PI3K-Glycolytic Pathway Regulation in Prostate Cancer. Clin Cancer Res, 2022. 28(16): p. 3603–3617.

30. Nguyen, B., et al., Pan-cancer Analysis of CDK12 Alterations Identifies a Subset of Prostate Cancers with Distinct Genomic and Clinical Characteristics. Eur Urol, 2020. 78(5): p. 671–679.

31. Hieronymus, H., et al., Copy number alteration burden predicts prostate cancer relapse. Proc Natl Acad Sci U S A, 2014. 111(30): p. 11139–44.

32. Ren, S., et al., Whole-genome and Transcriptome Sequencing of Prostate Cancer Identify New Genetic Alterations Driving Disease Progression. Eur Urol, 2018. 73(3): p. 322–339.

33. Weinstein, J.N., et al., The Cancer Genome Atlas Pan-Cancer analysis project. Nat Genet, 2013. 45(10): p. 1113–20.

34. Gao, D., et al., Organoid cultures derived from patients with advanced prostate cancer. Cell, 2014. 159(1): p. 176–187.

35. Tang, F., et al., Chromatin profiles classify castration-resistant prostate cancers suggesting therapeutic targets. Science, 2022. 376(6596): p. eabe1505.

36. Abida, W., et al., Prospective Genomic Profiling of Prostate Cancer Across Disease States Reveals Germline and Somatic Alterations That May Affect Clinical Decision Making. JCO Precis Oncol, 2017. 2017.

37. Sanjana, N.E., O. Shalem, and F. Zhang, Improved vectors and genome-wide libraries for CRISPR screening. Nature Methods, 2014. 11(8): p. 783–784.

38. McCabe, N., et al., BRCA2-deficient CAPAN-1 cells are extremely sensitive to the inhibition of Poly (ADP-Ribose) polymerase: an issue of potency. Cancer biology & therapy, 2005. 4(9): p. 934–936.

39. Blackford, A.N. and S.P. Jackson, ATM, ATR, and DNA-PK: The Trinity at the Heart of the DNA Damage Response. Molecular Cell, 2017. 66(6): p. 801–817.

40. Bass, T.E., et al., ATM Regulation of the Cohesin Complex Is Required for Repression of DNA Replication and Transcription in the Vicinity of DNA Double-Strand Breaks. Mol Cancer Res, 2023. 21(3): p. 261–273.

41. Kozlov, S.V., et al., Autophosphorylation and ATM activation: additional sites add to the complexity. J Biol Chem, 2011. 286(11): p. 9107–19.

42. Yap, T.A., et al., First-in-Human Trial of the Oral Ataxia Telangiectasia and RAD3-Related (ATR) Inhibitor BAY 1895344 in Patients with Advanced Solid Tumors. Cancer Discovery, 2021. 11(1): p. 80–91.

43. Yap, T.A., et al., Phase I Trial of First-in-Class ATR Inhibitor M6620 (VX-970) as Monotherapy or in Combination With Carboplatin in Patients With Advanced Solid Tumors. Journal of Clinical Oncology, 2020. 38(27): p. 3195–3204.

44. van Bussel, M.T.J., et al., A first-in-man phase 1 study of the DNA-dependent protein kinase inhibitor peposertib (formerly M3814) in patients with advanced solid tumours. Br J Cancer, 2021. 124(4): p. 728–735.

45. Zenke, F.T., et al., Pharmacologic Inhibitor of DNA-PK, M3814, Potentiates Radiotherapy and Regresses Human Tumors in Mouse Models. Molecular Cancer Therapeutics, 2020. 19(5): p. 1091–1101.

46. Gorecki, L., et al., Discovery of ATR kinase inhibitor berzosertib (VX-970, M6620): Clinical candidate for cancer therapy. Pharmacology & Therapeutics, 2020. 210: p. 107518.

47. Fokas, E., et al., Targeting ATR in vivo using the novel inhibitor VE-822 results in selective sensitization of pancreatic tumors to radiation. Cell Death Dis, 2012. 3(12): p. e441.

48. Wengner, A.M., et al., The Novel ATR Inhibitor BAY 1895344 Is Efficacious as Monotherapy and Combined with DNA Damage-Inducing or Repair-Compromising Therapies in Preclinical Cancer Models. Mol Cancer Ther, 2020. 19(1): p. 26–38.

49. Zhao, Y., et al., Preclinical Evaluation of a Potent Novel DNA-Dependent Protein Kinase Inhibitor NU7441. Cancer Research, 2006. 66(10): p. 5354–5362.

50. Brown, E.J. and D. Baltimore, Essential and dispensable roles of ATR in cell cycle arrest and genome maintenance. Genes Dev, 2003. 17(5): p. 615–28.

51. Caldecott, K.W., Causes and consequences of DNA single-strand breaks. Trends in Biochemical Sciences, 2024. 49(1): p. 68–78.

52. Izumi, N., et al., AAA+ proteins RUVBL1 and RUVBL2 coordinate PIKK activity and function in nonsense-mediated mRNA decay. Sci Signal, 2010. 3(116): p. ra27.

53. Neeb, A., et al., Advanced Prostate Cancer with ATM Loss: PARP and ATR Inhibitors. European Urology, 2021. 79(2): p. 200–211.

54. Tomimatsu, N., B. Mukherjee, and S. Burma, Distinct roles of ATR and DNA-PKcs in triggering DNA damage responses in ATM-deficient cells. EMBO Rep, 2009. 10(6): p. 629–35.

55. Finzel, A., et al., Hyperactivation of ATM upon DNA-PKcs inhibition modulates p53 dynamics and cell fate in response to DNA damage. Mol Biol Cell, 2016. 27(15): p. 2360–7.

56. Li, J. and D.F. Stern, Regulation of CHK2 by DNA-dependent protein kinase. J Biol Chem, 2005. 280(12): p. 12041–50.

57. Boehme, K.A., R. Kulikov, and C. Blattner, p53 stabilization in response to DNA damage requires Akt/PKB and DNA-PK. Proc Natl Acad Sci U S A, 2008. 105(22): p. 7785–90.

58. Callén, E., et al., Essential role for DNA-PKcs in DNA double-strand break repair and apoptosis in ATM-deficient lymphocytes. Mol Cell, 2009. 34(3): p. 285–97.

59. Schlam-Babayov, S., et al., Phosphoproteomics reveals novel modes of function and inter-relationships among PIKKs in response to genotoxic stress. The EMBO Journal, 2021. 40(2): p. e104400.

60. Yang, C.H., et al., MicroRNA203a suppresses glioma tumorigenesis through an ATM-dependent interferon response pathway. Oncotarget, 2017. 8(68): p. 112980–112991.

61. Cheng, L.C., et al., Phosphopeptide Enrichment Coupled with Label-free Quantitative Mass Spectrometry to Investigate the Phosphoproteome in Prostate Cancer. J Vis Exp, 2018(138).

62. Nita-Lazar, A., H. Saito-Benz, and F.M. White, Quantitative phosphoproteomics by mass spectrometry: past, present, and future. Proteomics, 2008. 8(21): p. 4433–43.

63. Hebert, A.S., et al., Comprehensive Single-Shot Proteomics with FAIMS on a Hybrid Orbitrap Mass Spectrometer. Anal Chem, 2018. 90(15): p. 9529–9537.

64. Swearingen, K.E. and R.L. Moritz, High-field asymmetric waveform ion mobility spectrometry for mass spectrometry-based proteomics. Expert Rev Proteomics, 2012. 9(5): p. 505–17.

65. Cox, J. and M. Mann, MaxQuant enables high peptide identification rates, individualized p.p.b.-range mass accuracies and proteome-wide protein quantification. Nat Biotechnol, 2008. 26(12): p. 1367–72.

66. Subramanian, A., et al., Gene set enrichment analysis: a knowledge-based approach for interpreting genome-wide expression profiles. Proc Natl Acad Sci U S A, 2005. 102(43): p. 15545–50.

67. Drake, J.M., et al., Phosphoproteome Integration Reveals Patient-Specific Networks in Prostate Cancer. Cell, 2016. 166(4): p. 1041–1054.

